# Genomic analysis reveals selection signatures of Yucatan miniature pig on its X chromosome during domestication

**DOI:** 10.1101/596387

**Authors:** Wei Zhang, Yuanlang Wang, Min Yang, Xudong Wu, Xiaodong Zhang, Yueyun Ding, Zongjun Yin

## Abstract

Yucatan miniature pig (YMP), a naturally small breed, has been domesticated in the hot and arid Yucatan Peninsula for a long time. However, its selection signatures on the X chromosome remain poorly understood. In this study, we focused on elucidating the selection signatures of YMP on the X chromosome during its domestication and breeding, using the whole-genome sequencing data. We performed population admixture analyses to determine its genetic relationships with other domesticated breeds and wild boars. Subsequently, we used two approaches, the fixation index (Fst) and π ratios, to identify the selection signatures with 100 kb windows sliding in 10 kb steps. As a result, we found that the ectodysplasin A (*EDA*) gene was related with hypoplasia or absence of hair and sweat glands. This could uncover the relative lack of odor in YMP and the presence of hypoplasia or absence of hair in pigs. Furthermore, we found several genes under selection in other animals. A bioinformatics analysis of the genes in selection regions showed that they were associated with growth, lipid metabolism, reproduction, and immune system. Our findings will lead to a better understanding of the unique genetic and phenotypic characteristics of YMP and offer a plausible method for their utilization as an animal model for hair and odor disease research.

## INTRODUCTION

Artificial selection plays a significant role in the domestication of livestock. The pig (*Sus scrofa domesticus*) was domesticated from *S. scrofa* approximately 9000 years ago in multiple regions of the world (Giuffra *et al*. 2000; Kijas and Andersson 2001; Larson *et al*. 2005). Compared with their wild counterparts, the domesticated animals showed notable changes in their adaptation strategies and phenotypic features. Detecting selection signatures in domesticated animals is important for elucidating the involvement of natural processes and human technology in the evolutionary process and how both have shaped modern animal genomes to provide insights for further improvement in livestock animals.

Theoretically, a novel beneficial variant undergoing selection usually shows a high population frequency and long-range linkage disequilibrium over a long period of time. (Grossman *et al*. 2010) Based on the decay of linkage disequilibrium and variation of allele frequency, a series of methods have been proposed (Sabeti PC, *et al*. 2002; Nielsen R. 2005; Voight BF, *et al*. 2006; Rubin C-J, *et al*. 2010; Lewontin *et al*. 1973; Tajima F 1983). These methods can be grouped into three categories: site-frequency spectrum, population differentiation, and linkage disequilibrium (Oleksyk *et al*. 2010). In recent years, many chip- or sequencing-based studies have found a series of genes associated with hair development, skin pigmentation, coat color, body size, fertility, environment adaptation and disease-resistance on humans (Sabeti *et al*. 2007; Fagny *et al*. 2014; Salas 2018) and a range of agricultural animals, including pig (Wang *et al*. 2014; Zhao *et al*. 2018; Moon *et al*. 2015), cattle (Kim *et al*. 2017; Pan *et al*. 2013; Qanbari *et al*., 2010), dog (Axelsson *et al*. 2013), goat and sheep (Alberto *et al*. 2018;Yang *et al*. 2016; Wang *et al*. 2016), chicken (Rubin *et al*. 2010;), and duck (Zhang *et al*. 2018). In addition, many studies have been performed by combine the Fst (Lewontin *et al*. 1973) and π (Tajima F. 1983) ratio approaches to detect selection signatures [Li *et al*. 2013, Lin *et al*. 2014; Zhang *et al*. 2018]. However, most of these studies focused on the autosomes. Evidence for the presence of selection signatures on their X chromosomes remains largely unexplored, and genes related to the traits of interest across domestic animals remain elusive. The pig X chromosome, approximately 125.94 Mb in size and accounting for 5.1% of the pig genome, contains 1172 genes, some of which are associated with growth, fat deposition, reproduction, disease, and other traits. Some examples of these genes are acyl-CoA synthetase long-chain family member 4 (*ACSL4*), ATPase Na+/K+ transporting family member beta 4 (*ATP1B4*), transient receptor potential channel 5 (*TRPC5*). Studies have shown that the X chromosome undergoes more drift than an autosome (Heyer and Segurel 2010), and the contributions of selection on the X chromosome and autosomes to the reduction in genome diversity have been estimated to range from 12 to 40% and 19 to 26%, respectively (McVicker *et al*. 2009). Therefore, it is necessary to investigate selection signatures on the X chromosome in pig to uncover its genetic characters. Rubin *et al*. (2012) pointed out that the X chromosome should be solely analyzed for the presence of selection signatures because it undergoes more drift than an autosome, and only sows should be used because they have different effective population sizes (Heyer and Segurel 2010).

Yucatan miniature pig (YMP), a naturally small breed, has been domesticated in the hot and arid Yucatan Peninsula and has distinct traits, such as gentleness, intelligence, disease resistance, and relative lack of odor (NRC 1991; Smith *et al*. 2005; Panepinto and Phillips 1986). Here, we downloaded the whole-genome sequences of 12 YMPs and 26 other wild and domesticated pigs. Subsequently, Fst and π ratio were implemented to identify selection signatures on the X chromosome of YMP. The findings herein will provide insights to increase understanding of the genetic base that determines the unique traits of YMP.

## MATERIALS AND METHODS

### Ethics statement

The entire procedure was carried out in strict accordance with the protocols approved by the Anhui Agricultural University Animal Ethics Committee under permission No. AHAU20140215.

### Experiment samples, mapping, and variants calling

Details about the sequenced samples can be found in previous studies (Zhao *et al*. 2018; Li *et al*. 2013; Kim *et al*. 2015). The sequencing data of 38 individuals were downloaded from the database of the National Center for Biotechnology Information (NCBI; https://www.ncbi.nlm.nih.gov/). These 38 individuals included 12 YMPs, 11 Asian wild boars, three European wild boars, and one each of the following pig breeds: Yorkshire, Landrace, Mini pig, Bamaxiang, Rongchang, Meishan, Tibetan, Laiwu, Wannan Black, *Sus barbatus*, *S. cebifrons*, *S. verrucosus*, and the common warthog *Phacochoerus africanus*. Before mapping, we removed the low-quality reads according to the method described by Zhao *et al*. (2018). The filtered reads were aligned to the reference pig genome (Sscrofa 11.1, GenBank assembly GCF_000003025.6) using BWA-MEM (Li and Durbin 2009). Subsequently, we used the open-source software packages Picard v1.119 (Picard, RRID: SCR_006525), GATK v3.0 (GATK, RRID: SCR_001876), and SAMtools v1.3.1 (SAMTOOLS, RRID: SCR_002105) (Li *et al*. 2009; McKenna *et al*. 2010) for downstream processing and variant calling. To filter out the variants and avoid possible false-positive results, the tool “VariantFiltration” of GATK was adopted with the following options: ““DP < 4.0 || DP > 1000.0 || MQ < 20.00” -cluster 2 -window 4” for single nucleotide variants (SNVs) and “QD < 2.0 || FS> 200.0 || ReadPosRankSum < −20.0” for insertions and deletions (InDels). After filtering, the variants were annotated using the ANNOVAR v2013–08-23 software (Wang *et al*. 2010).

### Phylogenetic tree construction, principal component analysis, and admixture analysis

To infer the genetic and population structure of pigs in our study, we converted the variant call format (VCF) file to the PLINK input files (.map and .ped). Firstly, we performed a principal component analysis (PCA) using the GCTA software (v1.25) (Yang *et al*. 2011) and employing “--make-grm” and “--grm tmp --pca 3” to generate the .eigenval and .eigenvec files. Secondly, we performed a population admixture analysis using the ADMIXTURE software v1.3 (ADMIXTURE, RRID: SCR_001263) (Holsinger and Weir 2009) in order to infer the true number of genetic populations (clusters or K) between the pig breeds. A cross-validation (CV) procedure were used to estimate the most preferable *K*-value, which exhibits a low cross-validation error compared to other *K*-values and takes the most probable number of inferred populations. Lastly, we performed a phylogenetic tree analysis by generating an identical by state (IBS) distance matrix through the PLINK software v1.90 (PLINK, RRID: SCR_001757) version (Purcell *et al*. 2007), constructed a neighbor-joining tree by SNPHYLO (Version: 20140701) (Lee *et al*. 2014) and drew a neighbor-joining tree using FigTree (v1.4.0) (Drummond *et al*. 2012).

### Detection of genome-wide selective sweeps

The regions under selection between the YMPs (n=12) and Asian wild boar (n=8) were identified based on two different statistics, i.e., Fst and π ratio. A 100 kb sliding window approach with 10 kb step-size was applied to calculate these statistics using the software PopGenome (Pfeifer *et al*. 2014). The genomic regions that overlapped between the two approaches were defined as selection signatures in YMP with simultaneously significantly high π ratio (1%) and Fst (1%) values.

### Functional enrichment analysis

To explore the potential biological significance of genes within these sweep regions, gene ontology (GO) and Kyoto Encyclopedia of Genes and Genomes (KEGG) analyses were carried out using the Database for Annotation, Visualization and Integrated Discovery (DAVID, v6.8) (Huang *et al*. 2009). The Benjamini-Hochberg FDR (false discovery rate) (Reiner *et al*. 2003) was used for correcting the P-values. Only the terms with P < 0.05 were considered as significant.

### Data Availability Statement

Illumina paired-end sequences for the pigs used in this study were downloaded from NCBI with accession numbers ERP001813, PRJNA260763 and SRA065461. All supporting data, File S1 contains detailed descriptions of selection signatures. File S2 contains selected genes within selection signatures. GO analysis were list in file S3. KEGG results were list in S4.

## RESULTS

### Genetic variations

After SNV and InDel calling, we identified 168 118 SNVs and 66 892 InDels (43 480 Inserts and 23 412 Deletions). Of all identified SNVs, the intergenic variants (67.7%) were the most common (Table 1). Meanwhile, 896 SNVs were synonymous and 359 were non-synonymous (Table 1). Of all identified InDels, the intergenic variants were the most common (67.9%). A total of 100 variants were found in the coding domain, and of the 80 frameshift variants, 55 were frameshift insertion and 25 were frameshift deletion (Table 1).

**Table 1.** Summary of identified SNVs and InDels.

| Variant type | Number of variants |
| --- | --- |
| SNV | 168 118 |
| Intergenic | 113 797 |
| Intron | 49 887 |
| Exon | 899 |
| Downstream | 1091 |
| Upstream | 978 |
| Splicing site | 11 |
| 5'-UTR | 306 |
| 3'-UTR | 1160 |
| Synonymous | 896 |
| Non-synonymous | 359 |
| InDel | 66 892 |
| Intergenic | 45 472 |
| Downstream | 440 |
| Upstream | 373 |
| Splicing site | 18 |
| 5'-UTR | 136 |
| 3'-UTR | 412 |
| Exon | 100 |
| Intron | 19 946 |
| Frameshift deletion | 25 |
| Frameshift insertion | 55 |
| Non-frameshift deletion | 12 |
| Non-frameshift insertion | 8 |

### Phylogenetic tree construction, PCA, and Admixture analyses

An unrooted phylogenetic tree was constructed to assess the phylogenetic relationship among the pig breeds (Fig. 1A). The results of the phylogenetic tree were grouped as expected and consistent with the PCA result of the present study (Fig. 1B), revealing strong clustering into three distinct genetic groups comprising Asian wild and domesticated pigs, European wild and domesticated pigs, and *Sus* species and *Phacochoerus africanus*. To infer population admixture, we first chose the lowest cross-validation error value (K=3, Fig. 1C), which was taken as the most probable number of inferred populations. Three clusters were observed: *P. africanus and Sus* species; European wild and domesticated pigs; Asian wild and domesticated pigs (Fig. 1D); these clusters showed a cluster pattern very similar to the PCA plots.

**Figure 1.**
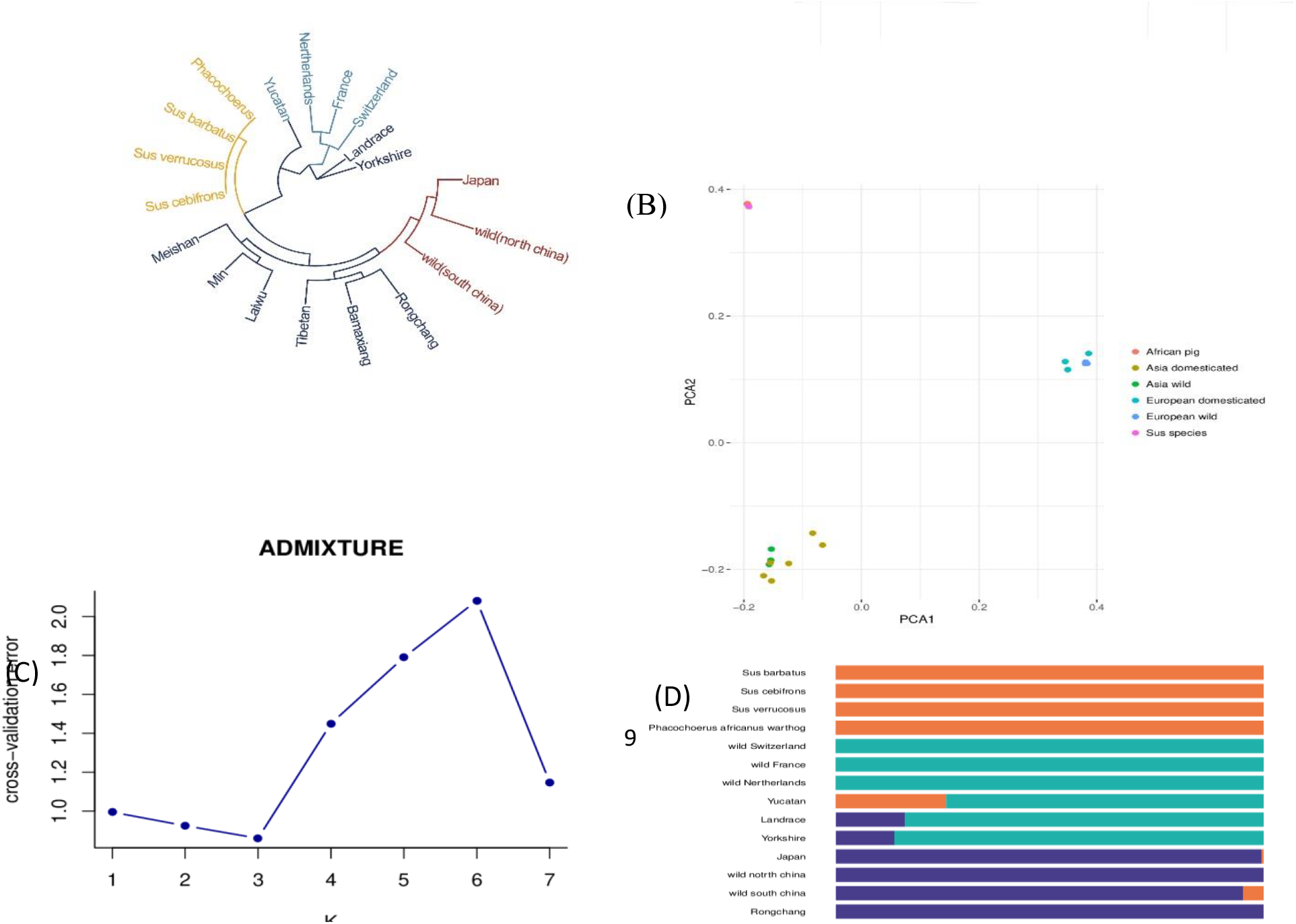
Population genetic structure of pig populations. (A) Neighbor-joining tree constructed from the single nucleotide variant (SNV) data of 19 pig subspecies. (B) Principal component analysis (PCA) plot of pig populations. Different colors represent different subspecies. (C) Cross-validation errors for diverse *k* values. As shown, *k* = 3 minimizes the cross-validation error. (D) Population structure of 19 pig populations. The lengths of different colors represent the proportions of ancestry from the ancestral populations. The names of pig breeds are indicated on the left.

### Selection detected by Fst and π ratio

The selection signatures of YMP on the X chromosome were identified based on the Fst and π ratio methods. The genome distributions of the two are shown in Fig 2A and 2B. A total of 316 selected regions (Additional file1: Table S1) were identified as having extremely high Fst values (1%) and significantly high π ratios (1%) (The Fst value was 0.761566, and the π ratio was 3.71527707). Seventy-seven genes were identified within the selected regions (Additional file1: Table S2). Further analysis of the identified genes by DAVID led to the identification of 34 categories of GO terms. The clusters were related to “reproductive process,” “signaling,” “immune system process,” “response to stimulus,” and “growth” (Fig 3A, Additional file1: Table S3). The KEGG analysis led to the enrichment of 62 pathways, most of which were related to immune system, lipid metabolism, digestive system, energy metabolism, cell growth and death, and signal transduction, for example, “protein digestion and absorption pathway,” “glycerolipid metabolism pathway,” “mTOR signaling pathway,” “oocyte meiosis pathway,” and “leukocyte transendothelial migration pathway”(Fig 3B, Additional file1: Table S4).

**Figure 2.**
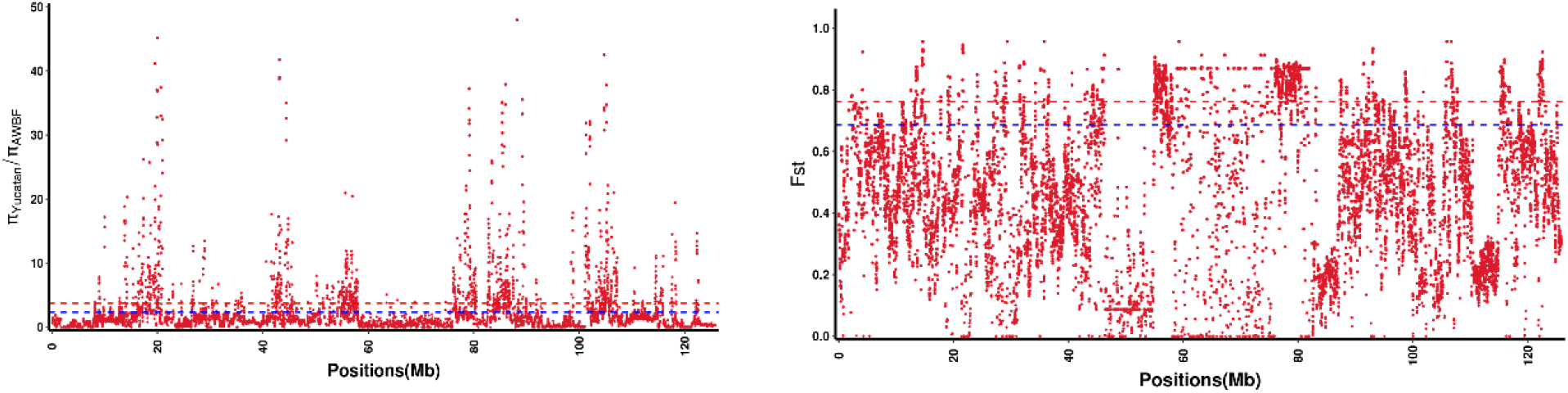
Distribution of Fst and π ratio calculated in 100 kb windows with 10 kb steps. (A) Distribution of π ratio; the red and blue lines represent the 0.01 and 0.05 levels, respectively. (B) Distribution of Fst; the red and blue lines represent the 0.01 and 0.05 levels, respectively.

**Fig. 3.**
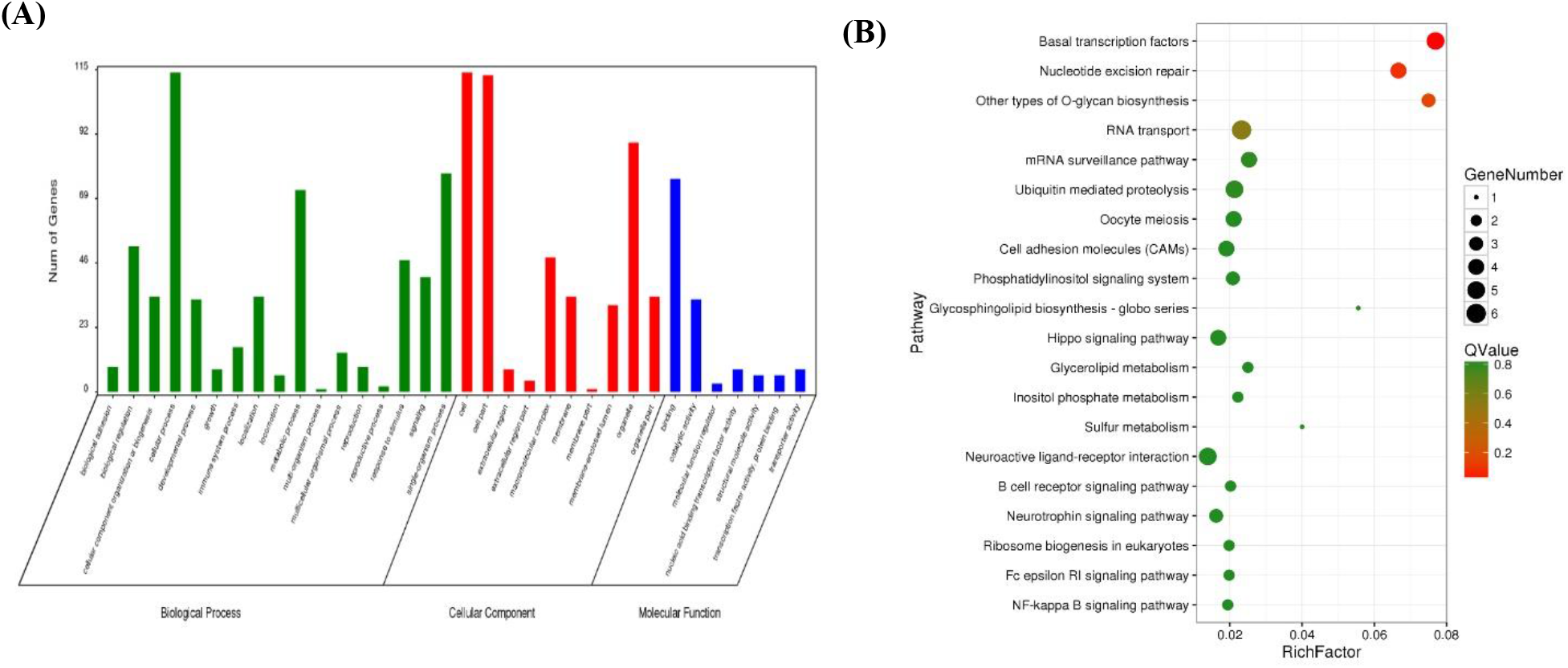
The Gene Ontology (GO) and Kyoto Encyclopedia of Genes and Genomes (KEGG) analysis of the genes harbored in the selection regions. (A) GO terms of the identified genes (B) Top 20 enriched pathways; the size of the dot indicates the number of genes, and the Q-value represents the difference level.

## DISCUSSION

Domesticated pig not only provides a great quantity of meat for humans, but is also used as a medical model for studying various diseases worldwide. To better understand the genetic basis of their domestication, we downloaded the whole-genome sequences of 12 YMPs and 26 pigs of other breeds from the NCBI databases. To our knowledge, the evidence for selection signatures on the X chromosome remains largely unexplored, and this is the first study to detect the selection signatures of YMP on its X chromosome. We observed 168 118 SNVs and 66 892 InDels in the YMP X chromosome and a low nucleotide diversity compared to its wild counterparts, suggesting artificial selection in YMP.

In recent years, selection signatures have been identified in humans and agricultural animals and crops, leading to the elucidation of the mechanisms of many complex traits and providing insight into the chip- or sequencing-based studies of various diseases. Considering more methods were combined to detect selection signatures, we employed two representative methods, Fst and π ratio, to explore YMP selection signatures on the X chromosome. Fst is effective for detecting selection footprints based on population differentiation. π ratio is a traditional and famous method. Ma *et al*. (2014) suggested cutoffs using the top 1% or 5% on the X chromosome to determine the significance of test statistic. Our results showed that a total of 317 overlapping selected regions were identified as having extremely high Fst values and significantly high π ratios. The regions around 55∼57 Mb and 75∼80 Mb were under strong selection according to the two approaches used in this study. Meanwhile, only small regions were detected around 46∼55 Mb and 58∼74 Mb; these regions might have been caused by X-inactivation resulting from sex-specific dosage compensation (SSDC). A total of 77 genes were harbored within the selected regions. The enrichment analysis of these genes revealed no significant term after correction, while six genes with P-values less than 0.05 indicated their biological information with the GO terms “Nucleotide excision repair”, Ubiquitin mediated proteolysis”, “Oocyte meiosis,” and “Cell adhesion molecules (CAMs).” Among these six genes, cullin 4B (*CUL4B*) plays a significant role in the development and progression of diffuse large B-cell lymphoma (*DLBCL*), while MiRNA-708 regulates the expression of *CUL4B* by binding to its 3′-UTR region (Li *et al*. 2019; Chen and Zhou 2018). The HECT, UBA, and WWE domain containing E3 ubiquitin protein ligase 1 (*HUWE1*) was suspected to be related with Say-Meyer syndrome and oxygen-glucose deprivation injury (Muthusamy *et al*. 2019). Studies on the structural maintenance of chromosome 1A (S*MC1A*) indicate that it plays an important role in development and behavior in Cornelia de Lange Syndrome (*CdLS*) (Mulder *et al*. 2019). Taking the above studies into consideration, we could conclude that YMP has suffered strong selection of different human diseases because of its use as a medical model.

Previous studies have shown that the porcine X chromosome harbors four quantitative trait loci (QTLs) strongly affecting backfat thickness (BFT), ham weight (HW), intramuscular fat content (IMF), and loin eye area (LEA); for example, acyl-CoA synthetase long-chain family member 4 (*ACSL4*), ATPase Na+/K+ transporting family member beta 4 (*ATP1B4*), and transient receptor potential channel 5 (*TRPC5*). In this study, we also identified the GO terms with lipid metabolism as “Glycerolipid metabolism” and “growth”. These two terms were related to five genes, including diacylglycerol kinase kappa (*DGKK*), ATRX chromatin remodeler (*ATRX*), mediator complex subunit 12 (*MED12*), TSPY-like 2 (*TSPYL2*), and myotubularin 1 (*MTM1*). ATRX is a chromatin-remodeling protein related to the alternative lengthening of telomeres (*ALT*), and it also affects other cellular functions related to epigenetic regulation (Hasse *et al*. 2018). MED12, a member of the Mediator kinase module, is an essential regulator of hematopoietic stem cell (HSC) homeostasis, is essential for early mouse development, regulates ovarian steroidogenesis, uterine development in mammalian egg, and plays an important role in neutrophil development in zebrafish (Aranda-Orgilles *et al*. 2016; Keightley *et al*. 2011; Wang *et al*. 2017; Rocha *et al*. 2010). TSPYL2 is a nucleosome assembly protein expressed in neuronal precursors and mature neurons. Studies have shown that disrupting the *Tspyl2* gene expression leads to behavioral and brain morphological alterations that mirror a number of neurodevelopmental psychiatric traits (Li *et al*. 2016). Mutations in *MTM1* result in X-linked myotubular/centronuclear myopathy (*XLMTM*) characterized by a severe decrease in muscle mass and strength in patients and murine models, leading to muscle hypotrophy (Bachmann *et al*. 2017; Al-Qusairi *et al*. 2013). DGKK, a master regulator, controls the switch between diacylglycerol and phosphatidic acid signaling pathways. A previous study has shown that DGKK controls synaptic protein translation and membrane properties by impacting lipid signaling in dendritic spine (Tabet *et al*. 2016).

Furthermore, reproduction-related genes were found to be selected in YMP. The mastermind-like domain-containing 1 gene (*MAMLD1*) encodes a mastermind-like domain-containing protein, which functions as a transcriptional co-activator. A previous study has shown that MAMLD1 contributes to the maintenance of postnatal testicular growth and daily sperm production (Miyado *et al*. 2017). Diaphanous-related formin 2 (*DIAPH2*) plays an important role in the gastrulation cell movement and development and normal function of ovaries. Defects in this gene have been linked to premature ovarian failure 2 (Lai *et al*. 2008; Bione *et al*. 1998). Testis expressed 11 (*TEX11*) is a target of the deleted in azoospermia-like (DAZL) present in the human foetal ovary. Studies have shown that *Dazl* deficiency in mouse results in an infertile phenotype in both sexes and has also been linked to infertility in humans (Rosario *et al*. 2017). *TEX11* has been identified as a selection gene in Australian Brahman cattle, which are associated with scrotal circumference (P < 0.001) and high percentage of normal sperm cells (P < 0.05), and could be used to select young bulls (Lyons *et al*. 2014). LDOC1 regulator of NFKB signaling (*LDOC1*), a paternally expressed gene, plays an important role in multinucleate syncytiotrophoblast formation and morphological diversification of placentas (Imakawa and Nakagawa 2017). A previous study has shown that the *Ldoc1* gene has undergone positive selection during eutherian evolution, which impacts reproductive fitness via the regulation of placental endocrine function (Naruse *et al*. 2014).

The immune system-related genes were also found to be selected in YMP. X-linked lymphoproliferative disease (XLP) is a primary immunodeficiency characterized by extreme susceptibility to Epstein-Barr virus, caused by a defect in the intracellular adapter protein SH2D1A. Studies have shown that the defective T and B cells exist in the absence of SH2D1A, likely explaining the progressive dysgammaglobulinemia in a subset of X-linked lympho-proliferative disease patients without the involvement of Epstein-Barr virus (Morra *et al*. 2005). Teneurin transmembrane protein 1 (*TENM1*) has been identified as a selected gene in dairy goat (Lai *et al*. 2016). It plays a role in congenital general anosmia and cerebral palsy. In *Drosophila melanogaster*, *TENM1* functions in synaptic-partner-matching between the axons of olfactory sensory neurons and target projection neurons and is involved in synapse organization in the olfactory system (Alkelai *et al*. 2016). Whole-exome sequencing has shown that *TENM1* act as a candidate gene for cerebral palsy (McMichael *et al*. 2015). The integral membrane protein 2a (Itm2a), one of the BRICHOS domain-containing proteins, plays a role in Dunnigan-type familial partial lipodystrophy (*FPLD2*) and regulates the development and function of T cells (Davies *et al*. 2017; Tai *et al*. 2014).

As is well known, YMP is associated with relative lack of odor. X-linked hypohidrotic ectodermal dysplasia (XLHED) is a genetic disease characterized by hypoplasia or absence of hair and sweat glands and is caused by variants in the *EDA* gene encoding ectodysplasin A (Pinheiro and Freire-Maia 1994; Kere *et al*. 1996). Studies have shown that variants in the *EDA* gene can lead to XLHED in humans, mice, and cattle (Cui and Schlessinger 2006). However, a frameshift variant (*c.842delT*) is a candidate causative variant for the observed XLHED in male puppies (Zhang *et al*. 2009). From the above studies, we could conclude that the trait of relative lack of odor in YMP is regulated by the *EDA* gene. Meanwhile, the *EDA* gene could explain the hypoplasia or absence of hair in pigs.

## CONCLUSION

In summary, our results elucidated the genetic relationship, PCA, and population admixture of YMP and pigs of other breeds from all over the world, using large amounts of sequencing data. To our knowledge, we detected the selection signatures of YMP on the X chromosome for the first time. Our analyses identified the positively selected genes, which were related to lipid metabolism, growth, immune system, and reproduction, in YMP. Moreover, we identified the *TEX11*, *LDOC1*, and *TENM1* genes under selection in YMP, as reported in other animals. The *EDA* gene was under selection and could explain the character of relative lack of odor in YMP and hypoplasia or absence of hair in pigs. Our data should be useful for better understanding of the mechanisms and identification of the targets of artificial selection in YMP.

## Acknowledgements

We thank Editage for English language editing.

---

| Chr | Start | End | Fst | Pi_Yucatan/Pi_AWBF |
| --- | --- | --- | --- | --- |
| X | 17720001 | 17820000 | 0.764167452 | 3.937198068 |
| X | 17730001 | 17830000 | 0.764167452 | 3.937198068 |
| X | 20860001 | 20960000 | 0.780876987 | 6.001976285 |
| X | 31450001 | 31550000 | 0.87368168 | 6.615942029 |
| X | 31460001 | 31560000 | 0.878633783 | 6.782608696 |
| X | 40590001 | 40690000 | 0.815366289 | 10.23188406 |
| X | 45000001 | 45100000 | 0.768483364 | 6.705163043 |
| X | 45010001 | 45110000 | 0.778219462 | 6.277173913 |
| X | 45020001 | 45120000 | 0.771116196 | 5.11548913 |
| X | 45930001 | 46030000 | 0.825677935 | 3.865665584 |
| X | 45940001 | 46040000 | 0.831247014 | 3.736589497 |
| X | 55080001 | 55180000 | 0.865218922 | 4.983016304 |
| X | 55090001 | 55190000 | 0.866007791 | 6.266983696 |
| X | 55100001 | 55200000 | 0.866303774 | 6.908967391 |
| X | 55110001 | 55210000 | 0.870513108 | 6.05298913 |
| X | 55120001 | 55220000 | 0.873838317 | 7.143342391 |
| X | 55160001 | 55260000 | 0.84795498 | 4.409267104 |
| X | 55170001 | 55270000 | 0.84795498 | 4.409267104 |
| X | 55180001 | 55280000 | 0.849211091 | 4.695642184 |
| X | 55190001 | 55290000 | 0.843456044 | 4.209259111 |
| X | 55250001 | 55350000 | 0.819509307 | 3.971014493 |
| X | 55300001 | 55400000 | 0.78442663 | 5.52415655 |
| X | 55310001 | 55410000 | 0.78442663 | 5.52415655 |
| X | 55320001 | 55420000 | 0.790013848 | 6.339373941 |
| X | 55350001 | 55450000 | 0.807816169 | 3.7671094 |
| X | 55360001 | 55460000 | 0.812384668 | 4.025607259 |
| X | 55390001 | 55490000 | 0.819013882 | 4.203595172 |
| X | 55560001 | 55660000 | 0.840571225 | 4.673007246 |
| X | 55570001 | 55670000 | 0.840571225 | 4.673007246 |
| X | 55580001 | 55680000 | 0.837840849 | 4.273550725 |
| X | 55590001 | 55690000 | 0.839427225 | 3.753623188 |
| X | 55600001 | 55700000 | 0.842252662 | 4.15307971 |
| X | 55620001 | 55720000 | 0.861228534 | 20.95953322 |
| X | 55630001 | 55730000 | 0.845965329 | 9.530370251 |
| X | 55640001 | 55740000 | 0.848291008 | 9.397385615 |
| X | 55650001 | 55750000 | 0.852730116 | 10.0073719 |
| X | 55660001 | 55760000 | 0.857408868 | 10.55395575 |
| X | 55670001 | 55770000 | 0.85643673 | 9.771347058 |
| X | 55680001 | 55780000 | 0.85643673 | 9.771347058 |
| X | 55690001 | 55790000 | 0.851260718 | 9.224763207 |
| X | 55700001 | 55800000 | 0.849510288 | 8.442154511 |
| X | 55710001 | 55810000 | 0.847389676 | 7.659545815 |
| X | 55720001 | 55820000 | 0.866928257 | 11.9236857 |
| X | 55750001 | 55850000 | 0.84294298 | 4.350845411 |
| X | 55770001 | 55870000 | 0.838514404 | 4.47705314 |
| X | 55780001 | 55880000 | 0.838514404 | 4.47705314 |
| X | 55790001 | 55890000 | 0.841197807 | 4.895531401 |
| X | 55800001 | 55900000 | 0.843454306 | 5.314009662 |
| X | 55810001 | 55910000 | 0.845378268 | 5.732487923 |
| X | 55820001 | 55920000 | 0.845378268 | 5.732487923 |
| X | 55830001 | 55930000 | 0.845378268 | 5.732487923 |
| X | 55840001 | 55940000 | 0.847038157 | 6.150966184 |
| X | 55850001 | 55950000 | 0.85198455 | 8.345788043 |
| X | 55860001 | 55960000 | 0.857466273 | 9.436141304 |
| X | 55870001 | 55970000 | 0.856594352 | 8.794157609 |
| X | 55880001 | 55980000 | 0.857466273 | 9.436141304 |
| X | 55890001 | 55990000 | 0.858900121 | 10.7201087 |
| X | 55900001 | 56000000 | 0.858228354 | 10.078125 |
| X | 55910001 | 56010000 | 0.857466273 | 9.436141304 |
| X | 55920001 | 56020000 | 0.85949673 | 11.36209239 |
| X | 55930001 | 56030000 | 0.858900121 | 10.7201087 |
| X | 55940001 | 56040000 | 0.858900121 | 10.7201087 |
| X | 56270001 | 56370000 | 0.815476461 | 3.759796336 |
| X | 56280001 | 56380000 | 0.815476461 | 3.759796336 |
| X | 56290001 | 56390000 | 0.824804034 | 5.510297483 |
| X | 56300001 | 56400000 | 0.821484012 | 5.611746758 |
| X | 56310001 | 56410000 | 0.833259382 | 6.466573165 |
| X | 56320001 | 56420000 | 0.837896829 | 9.04589372 |
| X | 56330001 | 56430000 | 0.826304421 | 4.863520347 |
| X | 56340001 | 56440000 | 0.825522059 | 4.265863213 |
| X | 56350001 | 56450000 | 0.825522059 | 4.265863213 |
| X | 56360001 | 56460000 | 0.825522059 | 4.265863213 |
| X | 56430001 | 56530000 | 0.844293478 | 4.449728261 |
| X | 56450001 | 56550000 | 0.847579452 | 4.877717391 |
| X | 56460001 | 56560000 | 0.851528231 | 4.748641304 |
| X | 56470001 | 56570000 | 0.854142287 | 5.604619565 |
| X | 56480001 | 56580000 | 0.856094557 | 6.460597826 |
| X | 56490001 | 56590000 | 0.859635756 | 6.75951087 |
| X | 56500001 | 56600000 | 0.859009197 | 6.331521739 |
| X | 56510001 | 56610000 | 0.852937371 | 5.176630435 |
| X | 56650001 | 56750000 | 0.836011281 | 4.533180778 |
| X | 56660001 | 56760000 | 0.841722177 | 4.768115942 |
| X | 56670001 | 56770000 | 0.841722177 | 4.768115942 |
| X | 56680001 | 56780000 | 0.8512593 | 5.579710145 |
| X | 56690001 | 56790000 | 0.83162924 | 4.320048309 |
| X | 56720001 | 56820000 | 0.846503162 | 3.939613527 |
| X | 56730001 | 56830000 | 0.851171144 | 5.004830918 |
| X | 56770001 | 56870000 | 0.852829364 | 11.20289855 |
| X | 56780001 | 56880000 | 0.852829364 | 11.20289855 |
| X | 56790001 | 56890000 | 0.852829364 | 11.20289855 |
| X | 56800001 | 56900000 | 0.857722108 | 11.84057971 |
| X | 56810001 | 56910000 | 0.855547636 | 10.01449275 |
| X | 56820001 | 56920000 | 0.855547636 | 10.01449275 |
| X | 56830001 | 56930000 | 0.860592137 | 10.65217391 |
| X | 56840001 | 56940000 | 0.862251656 | 7.666666667 |
| X | 56850001 | 56950000 | 0.862983884 | 8.579710145 |
| X | 56860001 | 56960000 | 0.864504045 | 11.31884058 |
| X | 56890001 | 56990000 | 0.86845381 | 20.44202899 |
| X | 56900001 | 57000000 | 0.806024402 | 4.707074429 |
| X | 56910001 | 57010000 | 0.810212873 | 4.97789241 |
| X | 56920001 | 57020000 | 0.806024402 | 4.707074429 |
| X | 56930001 | 57030000 | 0.791534664 | 4.247113731 |
| X | 56940001 | 57040000 | 0.807014061 | 5.059567674 |
| X | 57070001 | 57170000 | 0.839414231 | 4.082174619 |
| X | 57080001 | 57180000 | 0.83548518 | 4.185488425 |
| X | 57090001 | 57190000 | 0.831525176 | 3.757499294 |
| X | 57760001 | 57860000 | 0.78778284 | 8.956521739 |
| X | 57770001 | 57870000 | 0.78778284 | 8.956521739 |
| X | 57830001 | 57930000 | 0.840906022 | 4.692028986 |
| X | 57840001 | 57940000 | 0.83183691 | 5.605072464 |
| X | 57850001 | 57950000 | 0.8353898 | 6.137681159 |
| X | 57880001 | 57980000 | 0.836060411 | 6.876008729 |
| X | 57890001 | 57990000 | 0.841254139 | 7.178243028 |
| X | 63410001 | 63510000 | 0.78504553 | 5.072463768 |
| X | 68740001 | 68840000 | 0.798439242 | 3.913043478 |
| X | 68750001 | 68850000 | 0.798439242 | 3.913043478 |
| X | 68760001 | 68860000 | 0.798439242 | 3.913043478 |
| X | 74590001 | 74690000 | 0.778821879 | 3.897515528 |
| X | 74600001 | 74700000 | 0.778821879 | 3.897515528 |
| X | 74610001 | 74710000 | 0.778821879 | 3.897515528 |
| X | 76020001 | 76120000 | 0.845774688 | 5.537439614 |
| X | 76030001 | 76130000 | 0.849617774 | 6.602657005 |
| X | 76040001 | 76140000 | 0.854565878 | 6.974637681 |
| X | 76050001 | 76150000 | 0.851940994 | 5.90942029 |
| X | 76060001 | 76160000 | 0.851940994 | 5.90942029 |
| X | 76070001 | 76170000 | 0.839854699 | 7.625603865 |
| X | 76080001 | 76180000 | 0.831108144 | 6.027777778 |
| X | 76090001 | 76190000 | 0.839854699 | 7.625603865 |
| X | 76100001 | 76200000 | 0.845359883 | 9.223429952 |
| X | 76110001 | 76210000 | 0.843766428 | 8.690821256 |
| X | 76260001 | 76360000 | 0.860616423 | 5.822826087 |
| X | 76270001 | 76370000 | 0.846385312 | 5.547826087 |
| X | 76280001 | 76380000 | 0.874441133 | 7.425 |
| X | 76290001 | 76390000 | 0.874054739 | 8.17826087 |
| X | 76300001 | 76400000 | 0.865038532 | 5.643478261 |
| X | 76360001 | 76460000 | 0.862898171 | 6.264492754 |
| X | 76370001 | 76470000 | 0.866419885 | 6.103864734 |
| X | 76380001 | 76480000 | 0.842106059 | 5.309178744 |
| X | 76390001 | 76490000 | 0.838980626 | 4.776570048 |
| X | 76400001 | 76500000 | 0.838980626 | 4.776570048 |
| X | 76460001 | 76560000 | 0.854310559 | 4.311594203 |
| X | 76570001 | 76670000 | 0.81831686 | 3.937888199 |
| X | 76820001 | 76920000 | 0.843342105 | 4.419384058 |
| X | 76830001 | 76930000 | 0.864661159 | 13.89855072 |
| X | 76990001 | 77090000 | 0.852538956 | 5.537439614 |
| X | 77000001 | 77100000 | 0.85877014 | 6.442028986 |
| X | 77010001 | 77110000 | 0.857863354 | 5.90942029 |
| X | 77310001 | 77410000 | 0.842816851 | 5.492209338 |
| X | 77320001 | 77420000 | 0.842816851 | 5.492209338 |
| X | 77330001 | 77430000 | 0.850297325 | 7.599899472 |
| X | 77340001 | 77440000 | 0.83391774 | 5.277223773 |
| X | 77350001 | 77450000 | 0.848171153 | 10.95652174 |
| X | 77360001 | 77460000 | 0.848171153 | 10.95652174 |
| X | 77370001 | 77470000 | 0.851400445 | 12.7826087 |
| X | 77380001 | 77480000 | 0.848171153 | 10.95652174 |
| X | 77390001 | 77490000 | 0.848171153 | 10.95652174 |
| X | 77400001 | 77500000 | 0.852293032 | 10.68115942 |
| X | 77410001 | 77510000 | 0.857448325 | 11.31884058 |
| X | 77420001 | 77520000 | 0.857448325 | 11.31884058 |
| X | 77430001 | 77530000 | 0.856380309 | 10.4057971 |
| X | 77910001 | 78010000 | 0.828814229 | 4.88451087 |
| X | 77920001 | 78020000 | 0.843181818 | 5.46875 |
| X | 78280001 | 78380000 | 0.86627651 | 17.71014493 |
| X | 78290001 | 78390000 | 0.860800139 | 16.43478261 |
| X | 78300001 | 78400000 | 0.860800139 | 16.43478261 |
| X | 78310001 | 78410000 | 0.859697118 | 14.60869565 |
| X | 78320001 | 78420000 | 0.859697118 | 14.60869565 |
| X | 78330001 | 78430000 | 0.864865865 | 12.23188406 |
| X | 78340001 | 78440000 | 0.864865865 | 12.23188406 |
| X | 78350001 | 78450000 | 0.871142149 | 13.50724638 |
| X | 78360001 | 78460000 | 0.871366092 | 11.68115942 |
| X | 78370001 | 78470000 | 0.871664168 | 9.855072464 |
| X | 78490001 | 78590000 | 0.859697118 | 14.60869565 |
| X | 78500001 | 78600000 | 0.859697118 | 14.60869565 |
| X | 78510001 | 78610000 | 0.86627651 | 17.71014493 |
| X | 78520001 | 78620000 | 0.8668249 | 21.36231884 |
| X | 78530001 | 78630000 | 0.873957815 | 22.08959157 |
| X | 78540001 | 78640000 | 0.846056797 | 4.524587829 |
| X | 78550001 | 78650000 | 0.853200858 | 4.305238044 |
| X | 78800001 | 78900000 | 0.813479498 | 4.92599445 |
| X | 78810001 | 78910000 | 0.813479498 | 4.92599445 |
| X | 78820001 | 78920000 | 0.843478261 | 14.06521739 |
| X | 78830001 | 78930000 | 0.848150552 | 16.80434783 |
| X | 78840001 | 78940000 | 0.846044191 | 15.43478261 |
| X | 78850001 | 78950000 | 0.843478261 | 14.06521739 |
| X | 78860001 | 78960000 | 0.83619818 | 11.32608696 |
| X | 78870001 | 78970000 | 0.837747793 | 9.586956522 |
| X | 78900001 | 79000000 | 0.84311199 | 5.323242304 |
| X | 78910001 | 79010000 | 0.84311199 | 5.323242304 |
| X | 78920001 | 79020000 | 0.854684857 | 4.923593644 |
| X | 79020001 | 79120000 | 0.839890817 | 4.578804348 |
| X | 79030001 | 79130000 | 0.839890817 | 4.578804348 |
| X | 79040001 | 79140000 | 0.844593929 | 5.434782609 |
| X | 79080001 | 79180000 | 0.867834807 | 34.14492754 |
| X | 79090001 | 79190000 | 0.870167886 | 37.24637681 |
| X | 79100001 | 79200000 | 0.870167886 | 37.24637681 |
| X | 79110001 | 79210000 | 0.870268099 | 31.76811594 |
| X | 79120001 | 79220000 | 0.867373181 | 26.84057971 |
| X | 79130001 | 79230000 | 0.867738641 | 32.31884058 |
| X | 79140001 | 79240000 | 0.870309344 | 29.94202899 |
| X | 79150001 | 79250000 | 0.870468327 | 24.46376812 |
| X | 79160001 | 79260000 | 0.8707137 | 18.98550725 |
| X | 79170001 | 79270000 | 0.873715411 | 18.58272163 |
| X | 79200001 | 79300000 | 0.858222836 | 7.439285714 |
| X | 79210001 | 79310000 | 0.850763688 | 9.483850932 |
| X | 79220001 | 79320000 | 0.855416236 | 12.49689441 |
| X | 79230001 | 79330000 | 0.83857012 | 12.60450311 |
| X | 79240001 | 79340000 | 0.812836798 | 5.224976111 |
| X | 79250001 | 79350000 | 0.818519312 | 6.590778786 |
| X | 79260001 | 79360000 | 0.825533948 | 7.459925944 |
| X | 79270001 | 79370000 | 0.826119982 | 7.804347826 |
| X | 79280001 | 79380000 | 0.826119982 | 7.804347826 |
| X | 79290001 | 79390000 | 0.826119982 | 7.804347826 |
| X | 79300001 | 79400000 | 0.826776122 | 10.61141304 |
| X | 79310001 | 79410000 | 0.833076869 | 10.27513587 |
| X | 79320001 | 79420000 | 0.835597199 | 10.13315217 |
| X | 79330001 | 79430000 | 0.846999641 | 9.124320652 |
| X | 79430001 | 79530000 | 0.839004571 | 3.953125 |
| X | 79540001 | 79640000 | 0.8633752 | 6.797101449 |
| X | 79550001 | 79650000 | 0.862341479 | 5.731884058 |
| X | 79560001 | 79660000 | 0.862898171 | 6.264492754 |
| X | 79570001 | 79670000 | 0.8633752 | 6.797101449 |
| X | 79580001 | 79680000 | 0.8633752 | 6.797101449 |
| X | 79590001 | 79690000 | 0.8633752 | 6.797101449 |
| X | 79720001 | 79820000 | 0.851092171 | 6.367638965 |
| X | 79730001 | 79830000 | 0.846059148 | 5.778756192 |
| X | 79740001 | 79840000 | 0.838246864 | 4.40286186 |
| X | 79750001 | 79850000 | 0.841976322 | 4.298293891 |
| X | 79760001 | 79860000 | 0.845752236 | 4.991744634 |
| X | 79770001 | 79870000 | 0.845752236 | 4.991744634 |
| X | 79780001 | 79880000 | 0.848619008 | 5.685195377 |
| X | 79790001 | 79890000 | 0.848619008 | 5.685195377 |
| X | 79800001 | 79900000 | 0.844282983 | 5.443037975 |
| X | 79810001 | 79910000 | 0.844282983 | 5.443037975 |
| X | 79830001 | 79930000 | 0.855586999 | 8.152173913 |
| X | 79840001 | 79940000 | 0.834832456 | 4.263822115 |
| X | 80010001 | 80110000 | 0.841534242 | 3.745169082 |
| X | 80020001 | 80120000 | 0.861374169 | 8.555555556 |
| X | 80030001 | 80130000 | 0.860273292 | 7.490338164 |
| X | 80040001 | 80140000 | 0.860858391 | 8.02294686 |
| X | 80080001 | 80180000 | 0.842961261 | 6.923913043 |
| X | 80090001 | 80190000 | 0.836471978 | 5.360448808 |
| X | 80100001 | 80200000 | 0.84436756 | 6.060776064 |
| X | 80110001 | 80210000 | 0.842480263 | 5.648433848 |
| X | 80120001 | 80220000 | 0.830894592 | 3.999064984 |
| X | 80130001 | 80230000 | 0.830894592 | 3.999064984 |
| X | 80180001 | 80280000 | 0.845677725 | 4.193725922 |
| X | 80190001 | 80290000 | 0.850441852 | 4.748641304 |
| X | 80200001 | 80300000 | 0.844293478 | 4.449728261 |
| X | 80230001 | 80330000 | 0.85013906 | 3.922294172 |
| X | 80250001 | 80350000 | 0.872227152 | 8.913043478 |
| X | 80260001 | 80360000 | 0.867966373 | 8.275362319 |
| X | 80270001 | 80370000 | 0.868111252 | 9.188405797 |
| X | 80280001 | 80380000 | 0.868111252 | 9.188405797 |
| X | 80290001 | 80390000 | 0.868111252 | 9.188405797 |
| X | 80300001 | 80400000 | 0.868111252 | 9.188405797 |
| X | 80330001 | 80430000 | 0.826810924 | 4.200483092 |
| X | 80340001 | 80440000 | 0.834893506 | 5.11352657 |
| X | 80350001 | 80450000 | 0.8425533 | 6.483091787 |
| X | 80360001 | 80460000 | 0.837887773 | 5.570048309 |
| X | 80370001 | 80470000 | 0.83127409 | 4.657004831 |
| X | 80380001 | 80480000 | 0.83127409 | 4.657004831 |
| X | 80390001 | 80490000 | 0.83127409 | 4.657004831 |
| X | 80400001 | 80500000 | 0.826810924 | 4.200483092 |
| X | 80410001 | 80510000 | 0.810351967 | 3.97826087 |
| X | 87340001 | 87440000 | 0.830869029 | 7.79264214 |
| X | 87350001 | 87450000 | 0.790905605 | 6.599777035 |
| X | 87360001 | 87460000 | 0.768466915 | 6.109253066 |
| X | 87370001 | 87470000 | 0.783550853 | 6.811594203 |
| X | 87440001 | 87540000 | 0.81184152 | 12.7826087 |
| X | 87450001 | 87550000 | 0.832189169 | 10.95652174 |
| X | 87460001 | 87560000 | 0.81184152 | 12.7826087 |
| X | 87470001 | 87570000 | 0.798202165 | 10.95652174 |
| X | 87480001 | 87580000 | 0.823049174 | 9.130434783 |
| X | 87490001 | 87590000 | 0.807991958 | 7.304347826 |
| X | 91880001 | 91980000 | 0.770958889 | 4.213104715 |
| X | 98670001 | 98770000 | 0.831440287 | 17.07547525 |
| X | 98680001 | 98780000 | 0.831440287 | 17.07547525 |
| X | 98690001 | 98790000 | 0.831440287 | 17.07547525 |
| X | 98700001 | 98800000 | 0.831440287 | 17.07547525 |
| X | 98710001 | 98810000 | 0.827638366 | 15.10144928 |
| X | 98720001 | 98820000 | 0.827638366 | 15.10144928 |
| X | 98730001 | 98830000 | 0.822261689 | 17.85507246 |
| X | 98740001 | 98840000 | 0.806147881 | 11.68115942 |
| X | 98750001 | 98850000 | 0.800958836 | 4.608695652 |
| X | 98780001 | 98880000 | 0.798393797 | 6.913043478 |
| X | 98790001 | 98890000 | 0.795936203 | 9.152173913 |
| X | 98800001 | 98900000 | 0.792340292 | 7.217391304 |
| X | 98810001 | 98910000 | 0.786672085 | 8.782608696 |
| X | 101800001 | 101900000 | 0.792628736 | 7.623188406 |
| X | 106250001 | 106350000 | 0.840410883 | 12.28458498 |
| X | 106760001 | 106860000 | 0.866910056 | 4.811594203 |
| X | 106770001 | 106870000 | 0.818414322 | 4.144927536 |
| X | 106780001 | 106880000 | 0.820869565 | 6.463768116 |
| X | 106790001 | 106890000 | 0.811949853 | 9.217391304 |
| X | 106950001 | 107050000 | 0.775739377 | 8.876811594 |
| X | 107020001 | 107120000 | 0.768675111 | 6.532887402 |
| X | 107030001 | 107130000 | 0.766223833 | 5.473801561 |
| X | 115390001 | 115490000 | 0.778625722 | 4.862318841 |
| X | 115820001 | 115920000 | 0.811160865 | 3.736714976 |
| X | 115870001 | 115970000 | 0.887884058 | 9.510869565 |
| X | 115880001 | 115980000 | 0.887884058 | 9.510869565 |
| X | 115890001 | 115990000 | 0.897093238 | 11.10869565 |
| X | 115900001 | 116000000 | 0.897093238 | 11.10869565 |
| X | 122120001 | 122220000 | 0.864685969 | 4.184782609 |
| X | 122200001 | 122300000 | 0.890873098 | 4.162496597 |
| X | 122210001 | 122310000 | 0.889015147 | 3.844413358 |
| X | 122230001 | 122330000 | 0.880292165 | 11.67814794 |
| X | 122240001 | 122340000 | 0.829705278 | 12.7216262 |
| X | 122250001 | 122350000 | 0.838921589 | 14.66365519 |
| X | 122260001 | 122360000 | 0.838921589 | 14.66365519 |
| X | 122270001 | 122370000 | 0.825756453 | 13.38829287 |
| X | 122280001 | 122380000 | 0.838595036 | 12.11293055 |
| X | 122290001 | 122390000 | 0.763269909 | 11.44626388 |
| X | 122560001 | 122660000 | 0.899165569 | 7.31884058 |
| X | 122570001 | 122670000 | 0.893289435 | 6.985507246 |

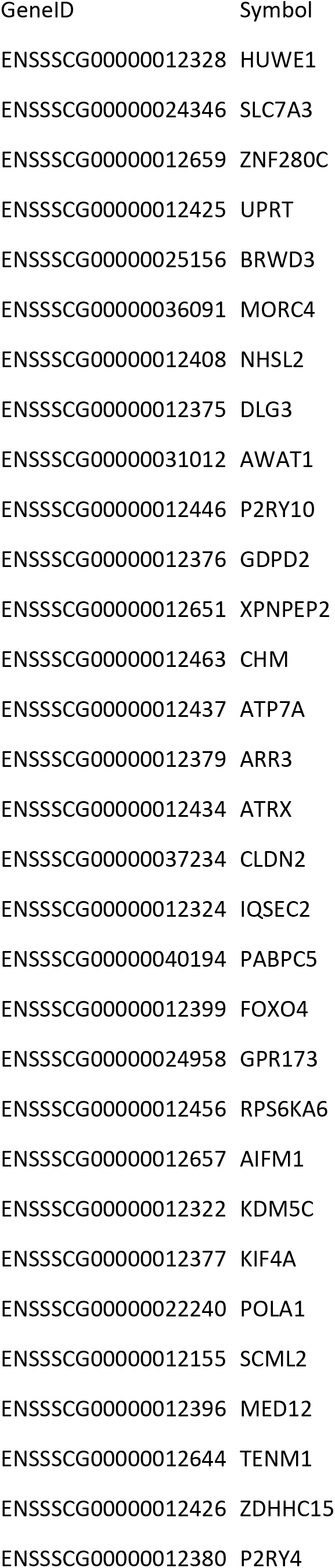
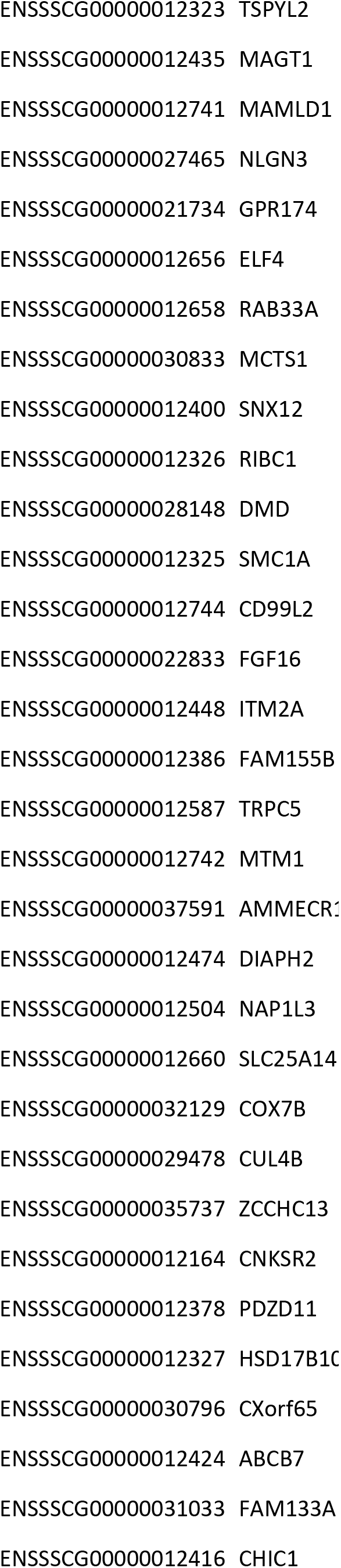
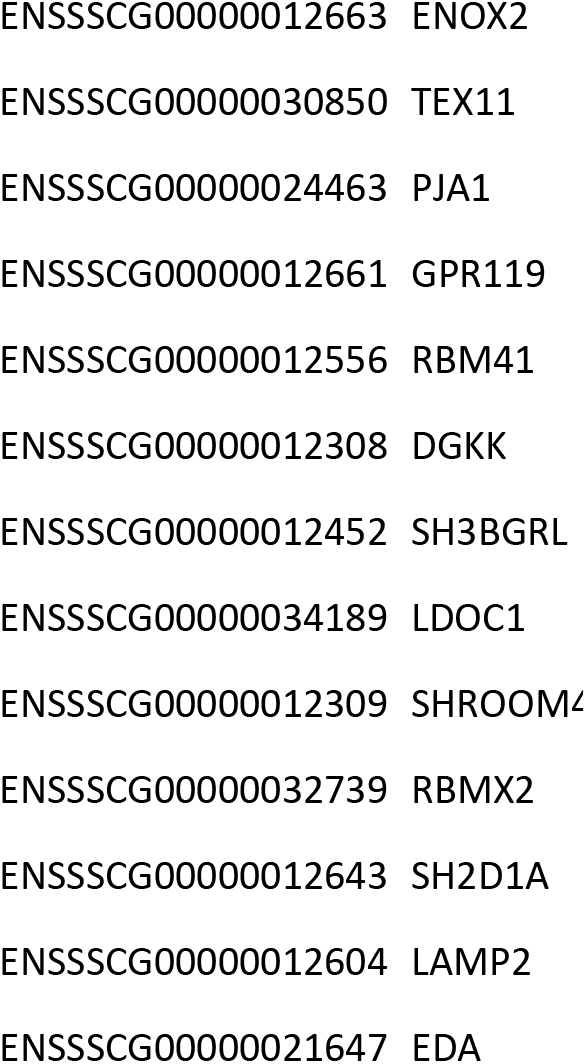

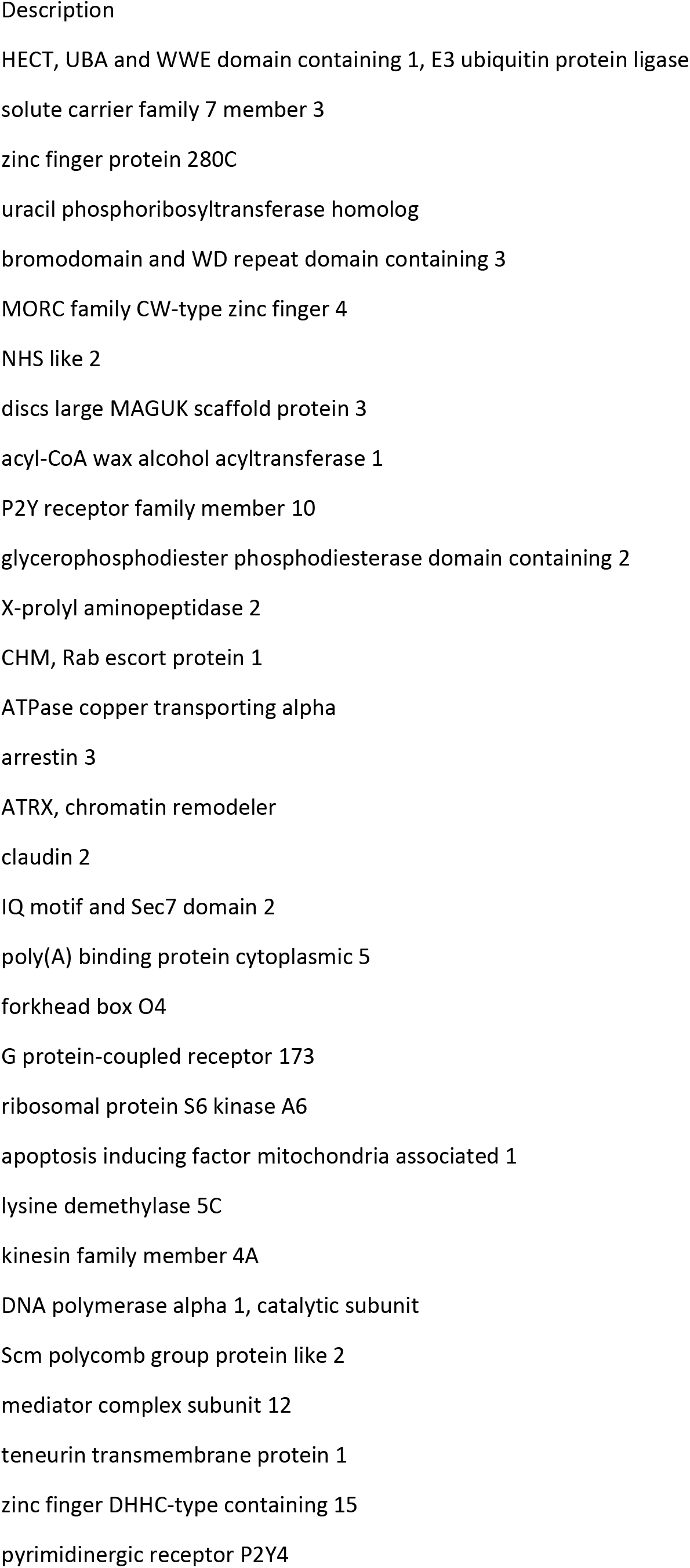
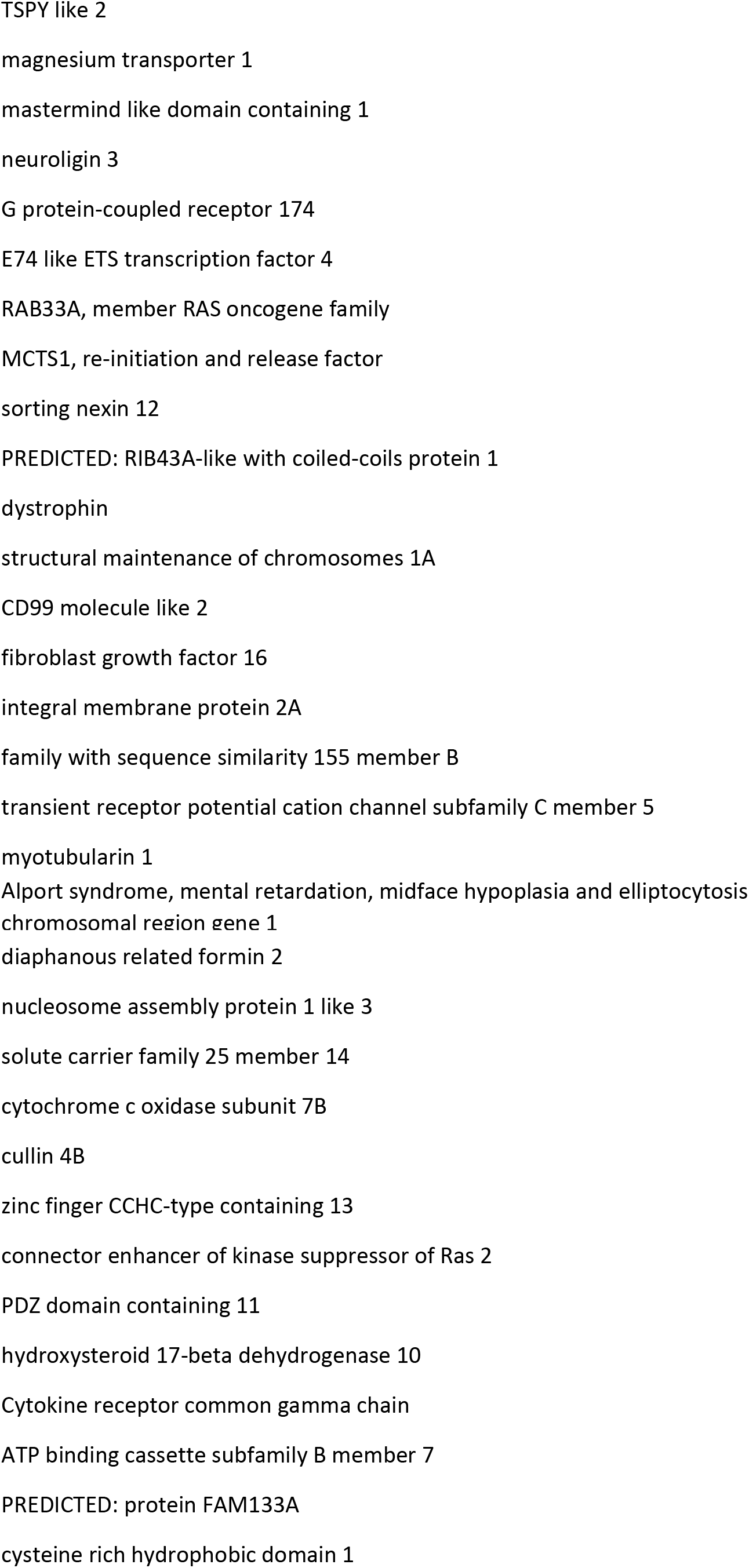
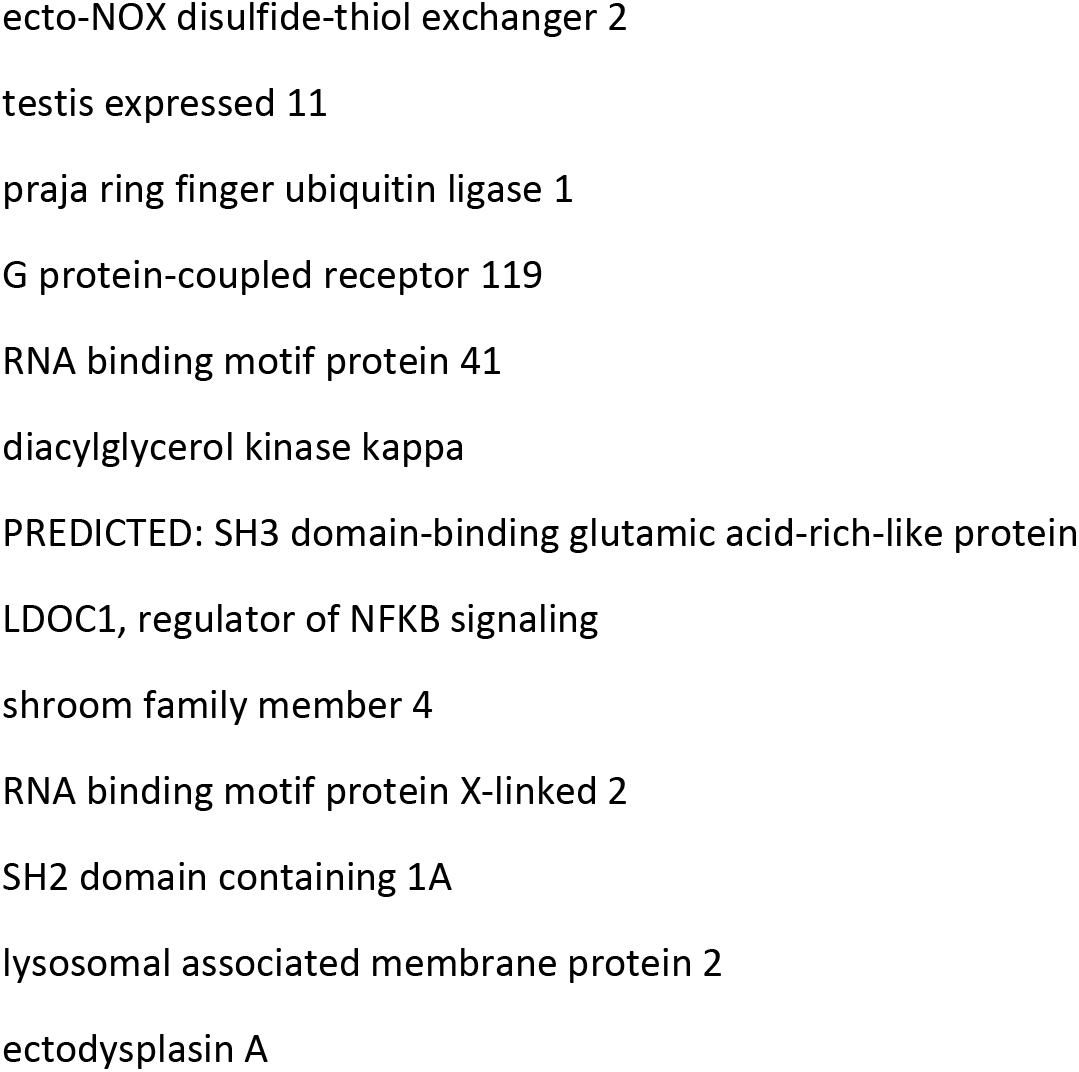

| Ontology | Class | Gene Number |
| --- | --- | --- |
| Biological Process | reproductive process | 2 |
| Biological Process | reproduction | 7 |
| Biological Process | cellular process | 78 |
| Biological Process | biological adhesion | 6 |
| Biological Process | growth | 4 |
| Biological Process | biological regulation | 35 |
| Biological Process | localization | 24 |
| Biological Process | developmental process | 22 |
| Biological Process | component organization or bic | 22 |
| Biological Process | immune system process | 8 |
| Biological Process | metabolic process | 46 |
| Biological Process | signaling | 26 |
| Biological Process | single-organism process | 54 |
| Biological Process | response to stimulus | 30 |
| Biological Process | multi-organism process | 1 |
| Biological Process | locomotion | 3 |
| Biological Process | multicellular organismal<br>process | 8 |
| Molecular Function | binding | 54 |
| Molecular Function | transcription factor activity,<br>protein binding | 2 |
| Molecular Function | transporter activity | 5 |
| Molecular Function | structural molecule activity | 4 |
| Molecular Function | nucleic acid binding<br>transcription factor activity | 4 |
| Molecular Function | catalytic activity | 21 |
| Molecular Function | molecular function regulator | 2 |
| Cellular Component | organelle | 60 |
| Cellular Component | macromolecular complex | 31 |
| Cellular Component | membrane-enclosed lumen | 19 |
| Cellular Component | cell part | 76 |
| Cellular Component | organelle part | 21 |
| Cellular Component | cell | 77 |
| Cellular Component | membrane | 22 |
| Cellular Component | membrane part | 1 |
| Cellular Component | extracellular region | 5 |
| Cellular Component | extracellular region part | 3 |

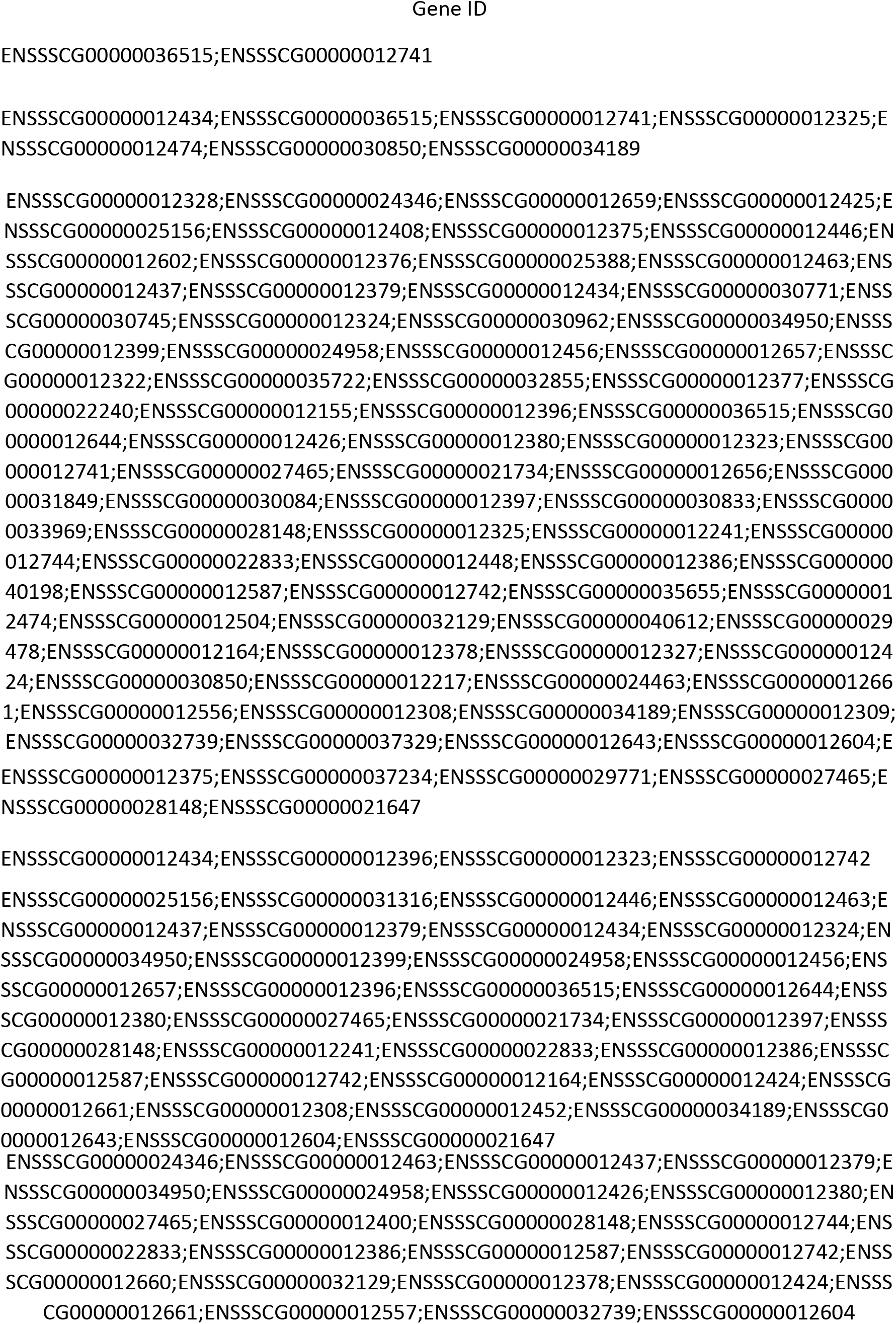
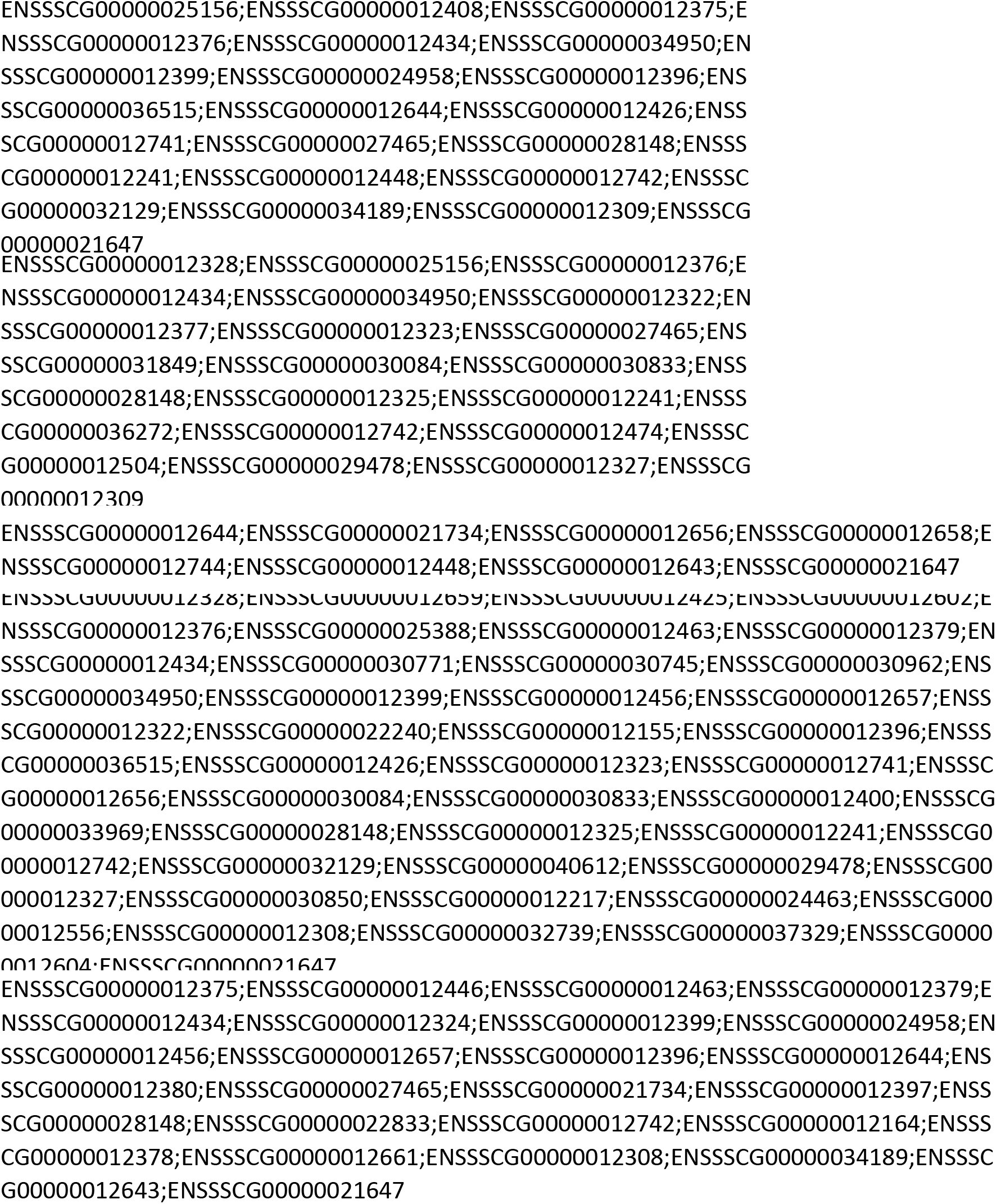
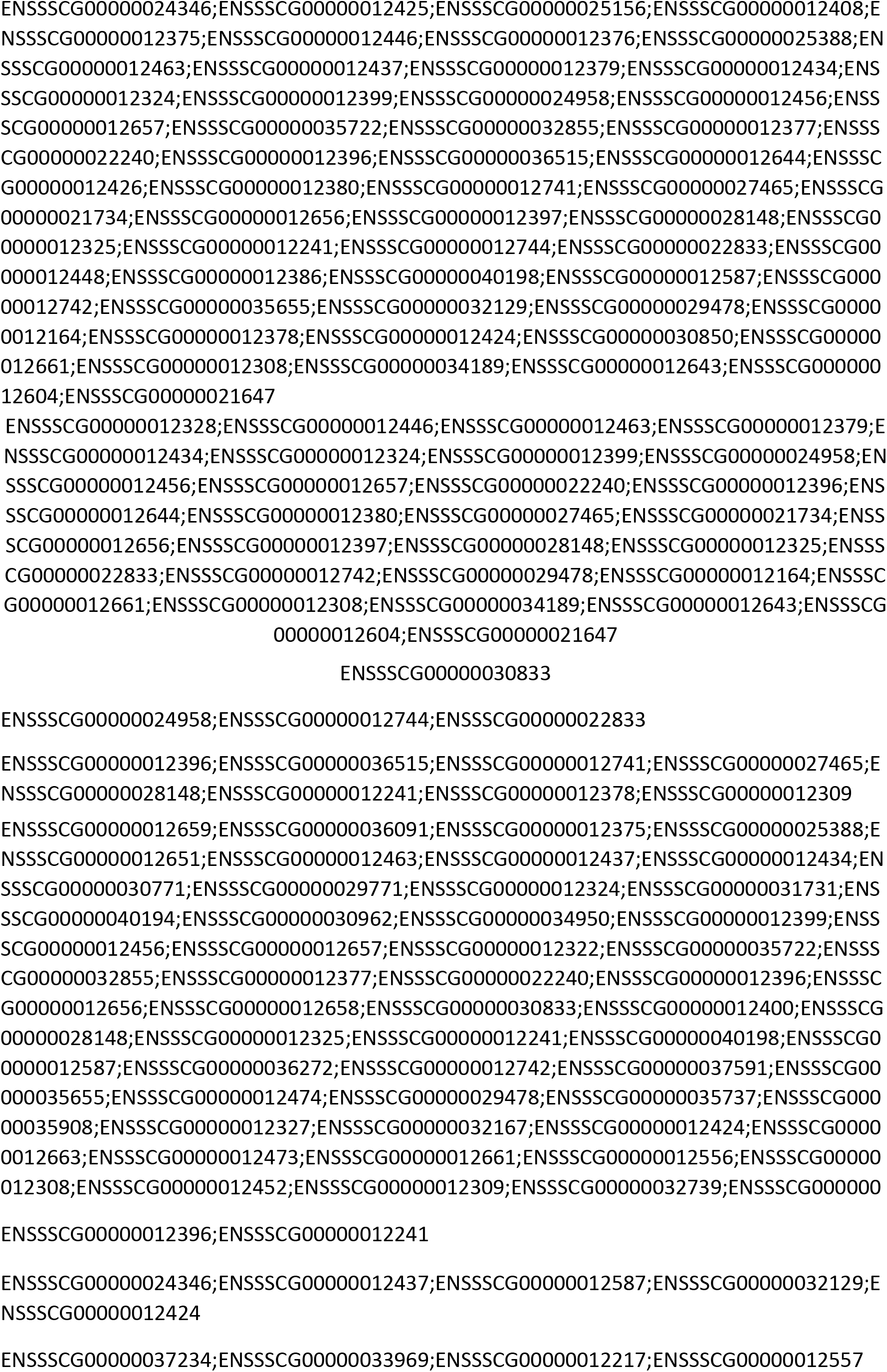
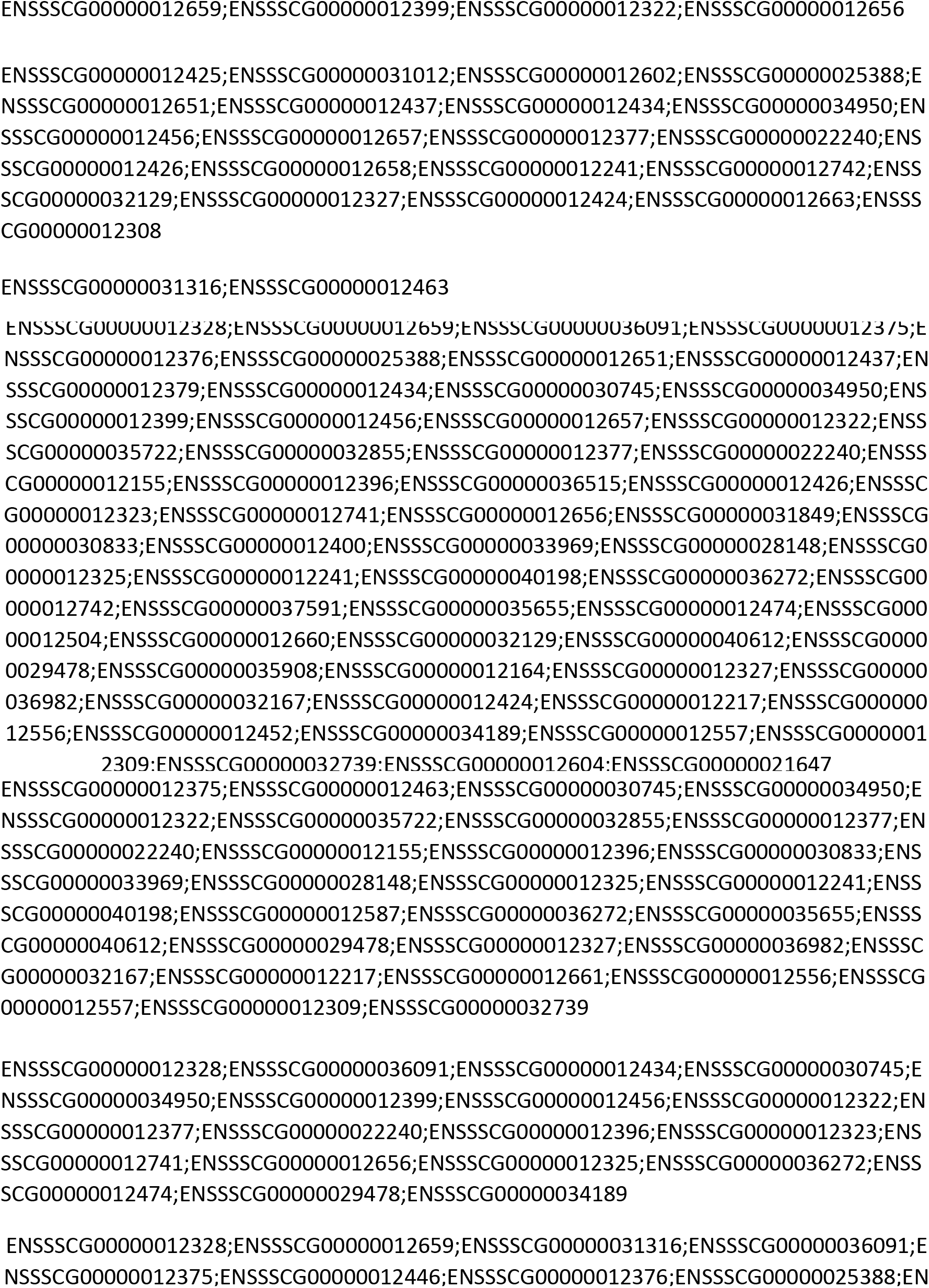
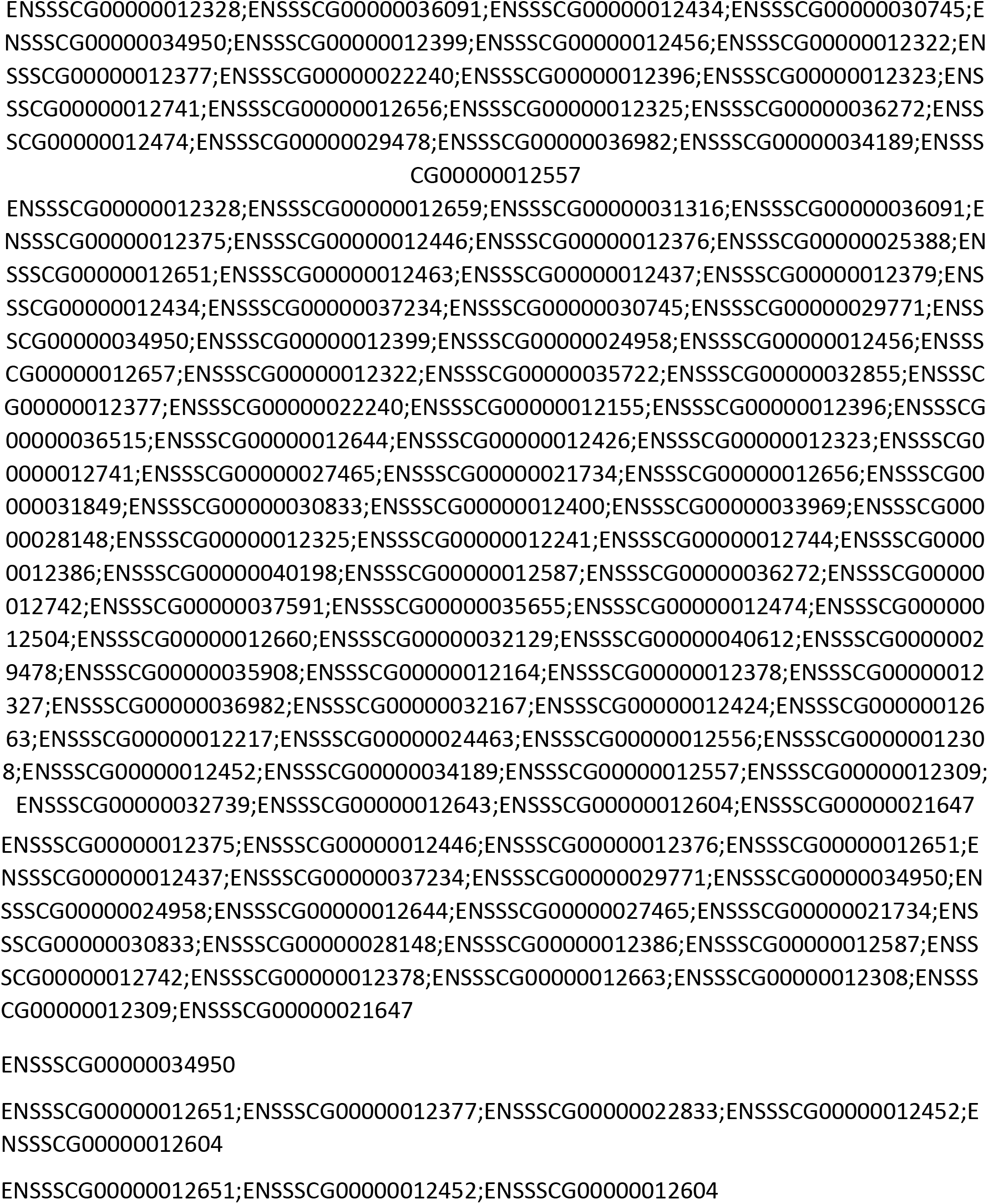

| KEGG_A_class | KEGG_B_class | Pathway | Pvalue | Qvalue |
| --- | --- | --- | --- | --- |
| Genetic Information Processing | Replication and repair | Nucleotide excision repair | 0.005523985 | 0.3424871 |
| Genetic Information Processing | Folding, sorting and degradation | Ubiquitin mediated proteolysis | 0.01304576 | 0.3896258 |
| Genetic Information Processing | Translation | RNA transport | 0.01885286 | 0.3896258 |
| Cellular Processes | Cell growth and death | Oocyte meiosis | 0.02697178 | 0.4180626 |
| Environmental Information Processing | Signaling molecules and interaction | Cell adhesion molecules (CAMs) | 0.03702157 | 0.4590675 |
| Environmental Information Processing | Signal transduction | Hippo signaling pathway | 0.05430543 | 0.5611561 |
| Metabolism | Energy metabolism | Sulfur metabolism | 0.1404866 | 0.727681 |
| Organismal Systems | Nervous system | Long-term potentiation | 0.1418819 | 0.727681 |
| Environmental Information Processing | Signal transduction | mTOR signaling pathway | 0.1481185 | 0.727681 |
| Cellular Processes | Cellular commiunity | Tight junction | 0.1547216 | 0.727681 |
| Cellular Processes | Cell growth and death | Apoptosis | 0.1565215 | 0.727681 |
| Genetic Information Processing | Folding, sorting and degradation | RNA degradation | 0.1692947 | 0.727681 |
| Metabolism | Nucleotide metabolism | Pyrimidine metabolism | 0.1844177 | 0.727681 |
| Organismal Systems | Endocrine system | Progesterone-mediated oocyte | 0.1997365 | 0.727681 |
| Environmental Information Processing | Signal transduction | Phosphatidylinositol signaling system | 0.2152102 | 0.727681 |
| Metabolism | Glycan biosynthesis and metabolism | Mucin type O-glycan biosynthesis | 0.2200569 | 0.727681 |
| Cellular Processes | Transport and catabolism | Endocytosis | 0.2237232 | 0.727681 |
| Metabolism | Metabolism of other amino acids | beta-Alanine metabolism | 0.2247832 | 0.727681 |
| Genetic Information Processing | Replication and repair | DNA replication | 0.2247832 | 0.727681 |
| Organismal Systems | Immune system | Natural killer cell mediated cytotoxicity | 0.2419885 | 0.727681 |
| Genetic Information Processing | Translation | mRNA surveillance pathway | 0.2464726 | 0.727681 |
| Cellular Processes | Cell growth and death | Cell cycle | 0.2846759 | 0.7401013 |
| Organismal Systems | Nervous system | Neurotrophin signaling pathway | 0.3071166 | 0.7401013 |
| Metabolism | Amino acid metabolism | Arginine and proline metabolism | 0.3219495 | 0.7401013 |
| Environmental Information Processing | Membrane transport | ABC transporters | 0.3219495 | 0.7401013 |
| Metabolism | Amino acid metabolism | Valine, leucine and isoleucine | 0.3260706 | 0.7401013 |
| Genetic Information Processing | Transcription | Basal transcription factors | 0.3260706 | 0.7401013 |
| Metabolism | Metabolism of other amino acids | Glutathione metabolism | 0.3342393 | 0.7401013 |
| Metabolism | Amino acid metabolism | Cysteine and methionine | 0.3581685 | 0.7459825 |
| Environmental Information Processing | Signal transduction | MAPK signaling pathway | 0.3803863 | 0.7459825 |
| Genetic Information Processing | Transcription | Spliceosome | 0.3822052 | 0.7459825 |
| Metabolism | Lipid metabolism | Glycerolipid metabolism | 0.3850232 | 0.7459825 |
| Metabolism | Carbohydrate metabolism | Inositol phosphate metabolism | 0.4214888 | 0.7685972 |
| Organismal Systems | Sensory system | Taste transduction | 0.4214888 | 0.7685972 |
| Cellular Processes | Cell growth and death | p53 signaling pathway | 0.4524925 | 0.7907725 |
| Genetic Information Processing | Translation | Ribosome biogenesis in eukaryotes | 0.4591582 | 0.7907725 |
| Organismal Systems | Digestive system | Protein digestion and absorption | 0.5036442 | 0.8276763 |
| Environmental Information Processing | Signaling molecules and interaction | Cytokine-cytokine receptor interaction | 0.5132722 | 0.8276763 |
| Organismal Systems | Endocrine system | Insulin secretion | 0.5215943 | 0.8276763 |
| Organismal Systems | Circulatory system | Cardiac muscle contraction | 0.5389086 | 0.8276763 |
| Metabolism | Lipid metabolism | Glycerophospholipid metabolism | 0.5473343 | 0.8276763 |
| Environmental Information Processing | Signal transduction | Ras signaling pathway | 0.5719371 | 0.8296981 |
| Cellular Processes | Cell motility | Regulation of actin cytoskeleton | 0.5754358 | 0.8296981 |
| Organismal Systems | Immune system | Leukocyte transendothelial | 0.6143165 | 0.8475808 |
| Cellular Processes | Transport and catabolism | Lysosome | 0.619047 | 0.8475808 |
| Organismal Systems | Endocrine system | Thyroid hormone signaling pathway | 0.6329001 | 0.8475808 |
| Environmental Information Processing | Signaling molecules and interaction | Neuroactive ligand-receptor interaction | 0.6457787 | 0.8475808 |
| Genetic Information Processing | Translation | Ribosome | 0.6735485 | 0.8475808 |
| Cellular Processes | Transport and catabolism | Phagosome | 0.70072 | 0.8475808 |
| Metabolism | Global and Overview | Metabolic pathways | 0.7026534 | 0.8475808 |
| Environmental Information Processing | Signal transduction | Jak-STAT signaling pathway | 0.7044104 | 0.8475808 |
| Environmental Information Processing | Signal transduction | FoxO signaling pathway | 0.7222047 | 0.8475808 |
| Environmental Information Processing | Signal transduction | Phospholipase D signaling pathway | 0.7323734 | 0.8475808 |
| Metabolism | Energy metabolism | Oxidative phosphorylation | 0.7531719 | 0.8475808 |
| Metabolism | Global and Overview | Microbial metabolism in diverse | 0.7607428 | 0.8475808 |
| Genetic Information Processing | Folding, sorting and degradation | Protein processing in endoplasmic | 0.7695282 | 0.8475808 |
| Environmental Information Processing | Signal transduction | PI3K-Akt signaling pathway | 0.7848277 | 0.8475808 |
| Metabolism | Nucleotide metabolism | Purine metabolism | 0.7940326 | 0.8475808 |
| Metabolism | Global and Overview | Biosynthesis of secondary | 0.817143 | 0.8475808 |
| Metabolism | Global and Overview | Biosynthesis of antibiotics | 0.8417184 | 0.8475808 |
| Environmental Information Processing | Signal transduction | Rap1 signaling pathway | 0.8436966 | 0.8475808 |
| Environmental Information Processing | Signal transduction | cAMP signaling pathway | 0.8475808 | 0.8475808 |

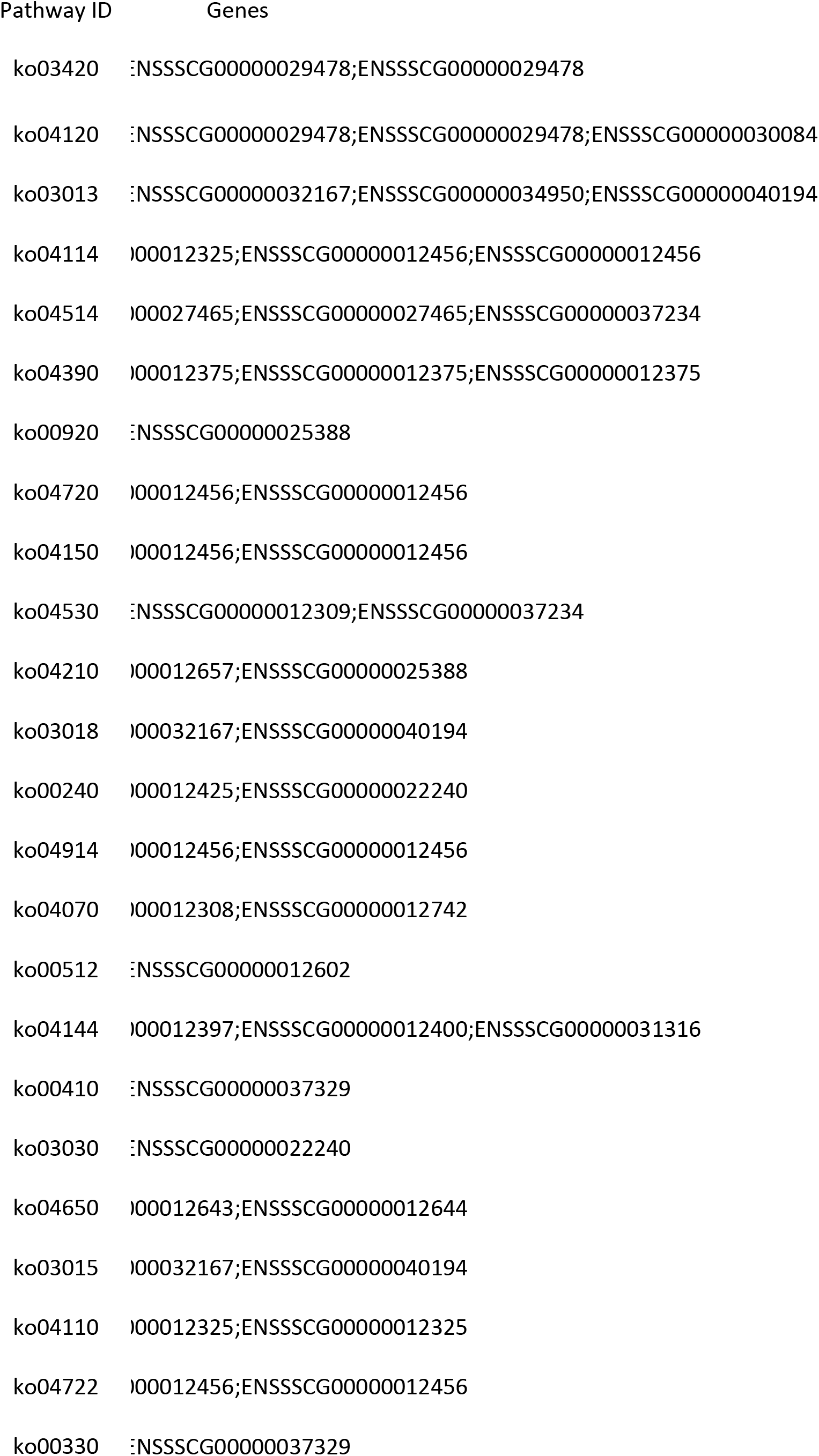
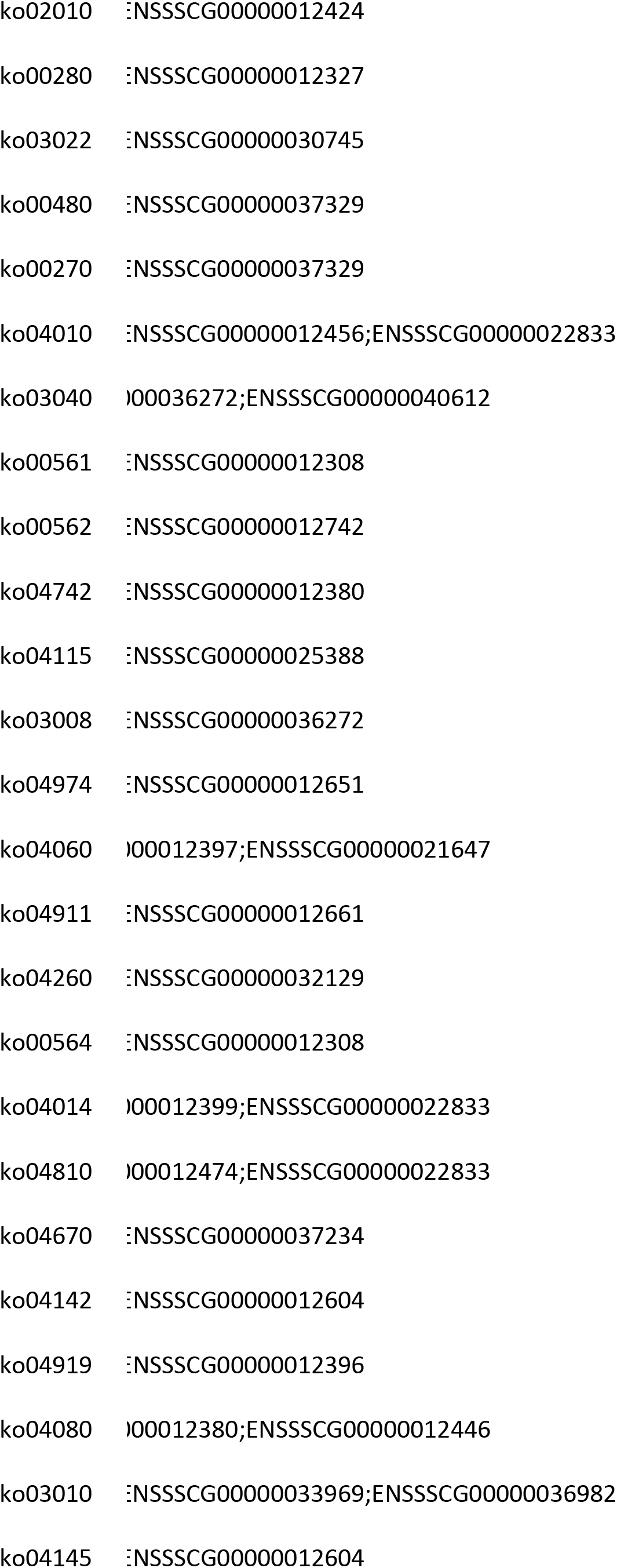
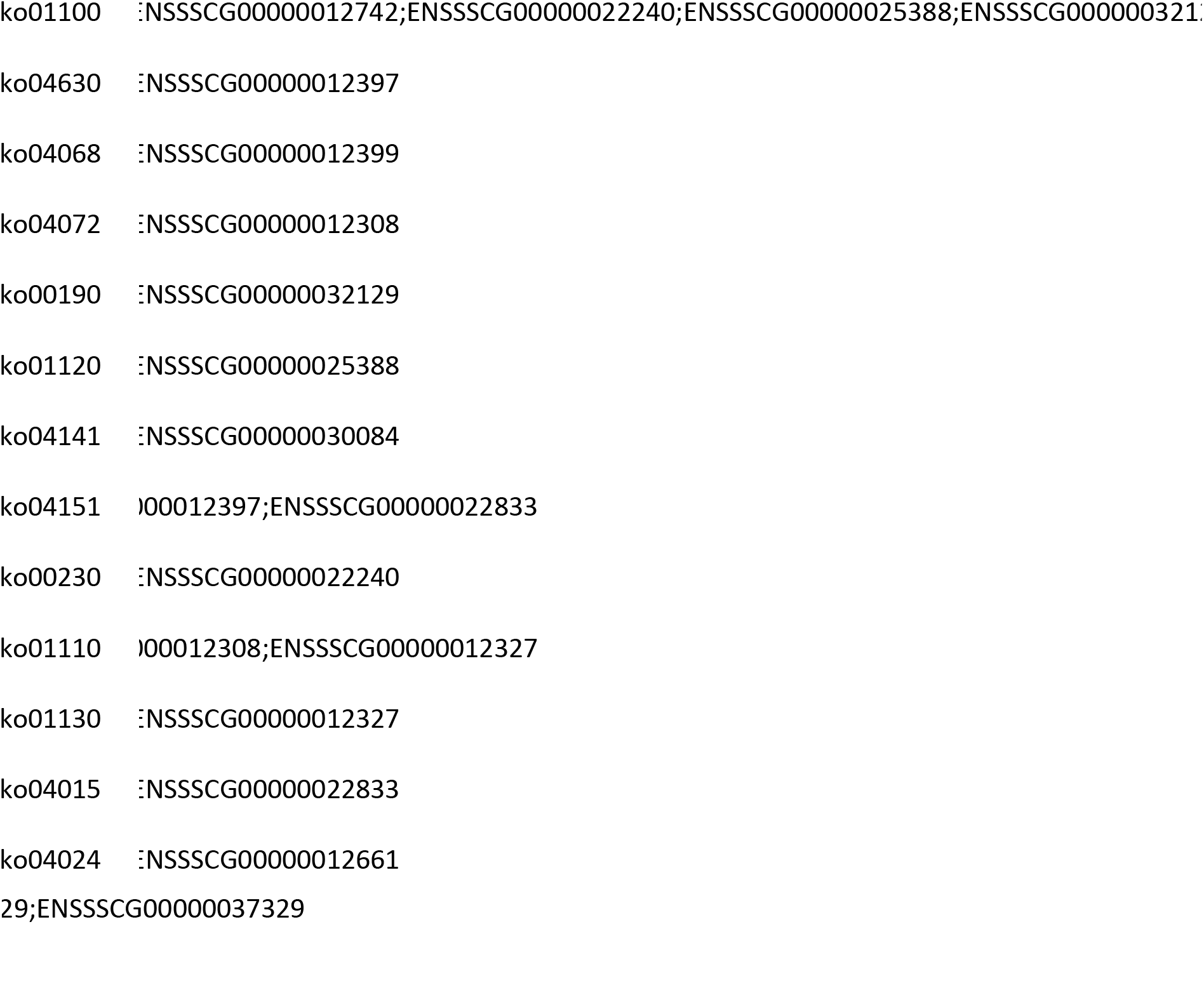

